# A biologically-informed polygenic score identifies endophenotypes and clinical conditions associated with the insulin receptor function on specific brain regions

**DOI:** 10.1101/289983

**Authors:** Shantala A. Hari Dass, Kathryn McCracken, Irina Pokhvisneva, Lawrence M. Chen, Elika Garg, Thao T. T. Nguyen, Zihan Wang, Barbara Barth, Moein Yaqubi, Lisa M. McEwen, Julie L. MacIsaac, Josie Diorio, Michael S. Kobor, Kieran J. O’Donnell, Michael J. Meaney, Patricia P. Silveira

## Abstract

Conventional polygenic scores derived from genome-wide association studies do not reflect gene networks that code for biological functions. We present an alternative approach creating a biologically-informed polygenic score based on the insulin receptor (IR) gene networks in the mesocorticolimbic system and hippocampus that regulate reward sensitivity/inhibitory control and memory, respectively. Across multiple samples (n = 4300) our biologically-informed IR-PRS score showed better prediction of child impulsivity and cognitive performance, as well as risk for early addiction onset and Alzheimer’s disease in comparison to conventional polygenic scores for ADHD, addiction and dementia. This novel, biologically-informed approach enables the use of genomic datasets to probe relevant biological processes involved in neural function and disorders.

**One Sentence Summary:** A polygenic score based on genes co-expressed with the insulin receptor predicts childhood behavior and adult disease.

## Main Text

The co-morbidity between metabolic and neuropsychiatric disorders is well-established, but poorly understood. The high co-occurrence of several psychiatric conditions (e.g. major depression, bipolar disorder, dementia) with insulin resistance suggests a common underlying mechanism (Anderson, Freedland et al. 2001). Insulin receptors (IR) are expressed throughout the brain, in areas including the ventral tegmental area, striatum, prefrontal cortex and hippocampus. Insulin is actively transported across the blood-brain barrier, and its action on mesocorticolimbic receptors modulates synaptic plasticity in dopaminergic neurons, affecting dopamine-related behaviors such as response to reward, impulsivity and decision-making (Kullmann, Heni et al. 2016). IR location on hippocampal glutamatergic synapses suggests a role of insulin in neurotransmission, synaptic plasticity and modulation of learning and memory, while its inhibition is described in Alzheimer’s disease and related animal models (Bomfim, Forny-Germano et al. 2012).

Genetic studies can be a pertinent tool to investigate the neurobiological mechanisms that explain the co-morbidity between metabolic and neuropsychiatric conditions. Genome-wide association studies (GWAS) provide the basis for cumulative variants that associate with health outcomes and reflect genetic predispositions to common disorders where individual variants carry small effects. The cumulative polygenetic risk of the individual can be used to estimate risk through polygenic risk scores (PRS), by multiplying the measured number of risk alleles at a locus by the effect size of the association between a particular genotype and the outcome from the relevant GWAS study, and summing across the genome (Wray and Goddard 2010). GWAS and PRS methodologies are focused on statistically significant candidate associations between scattered loci and a certain condition or trait, not accounting for the fact that genes operate in networks, and code for precise biological functions in specific tissues. Although recent studies explore the ability of conventional polygenic risk scores (PRS) to detect vulnerability endophenotypes, the conservative approach does not provide information about the underlying network in which these genes operate, nor the mechanism of the disease. This makes the biological underpinnings of the PRS complicated to disentangle, restricting their clinical scope/implications. Since the PRS are conventionally based on statistical thresholds, while they might associate with the final disease state, they are not informative of the various clinical manifestations of the disease, and endophenotypes that may precede the disease development. A score that captures a genetic risk to a clinical manifestation-potentially an early onset manifestation-would be able to mold subsequent clinical and therapeutic endeavors. With the reduction in cost of genotyping strategies we are poised to expand the implications of a multitude of GWA studies into healthcare. Hence, we aimed to address this need by developing a novel genomics approach that provides a biologically-informed genetic score, based on genes co-expressed with the IR in specific brain regions. Our new methodology enabled a hypothesis-driven, neurobiological analysis of phenotypes linked to brain disorders across developmental stages. It allowed us to identify differences in dopamine-related behavioral outcomes, namely impulsivity and risk for addiction, when focused on the mesocorticolimbic system. By comparison, the hippocampal IR gene network permitted the distinction of differences in childhood cognitive performance, and the presence of Alzheimer’s disease in adults.

### Establishment of the mesocorticolimbic biologically-informed genetic score based on genes co-expressed with the insulin receptor (ePRS-IR)

The steps for the development of the IR expression-based polygenic risk score (ePRS-IR) are depicted in Figure 1 (Fig. 1A). We (a) extracted brain-region specific IR co-expression data from publicly available RNA sequencing data sets in mice (Mulligan, Mozhui et al. 2017); (b) generated a co-expression matrix with IR in the ventral striatum and in the prefrontal cortex using transcriptomic data (i.e., the mesolimbic IR matrix), only including genes with a correlation coefficient >|0.5|; (c) identified consensus autosomal transcripts from this gene list, performing a developmental enrichment within the same brain areas, by selecting transcripts differentially expressed at ≥1.5 fold during fetal and child development as compared to adult samples (Miller, Ding et al. 2014); the final list included 281 genes; (d) based on their functional annotation, we gathered all the existing SNPs from these genes (total = 11,068); (e) subjected this list of SNPs to linkage disequilibrium clumping (Wray, Lee et al. 2014), to inform removal of highly correlated SNPs, resulting in 1897 independent functional SNPs and (f) compiled these SNPs in a polygenic score(Wray and Goddard 2010) using the association betas described in a conventional ADHD GWAS (Neale, Medland et al. 2010) (Table S1).

**Fig. 1.**
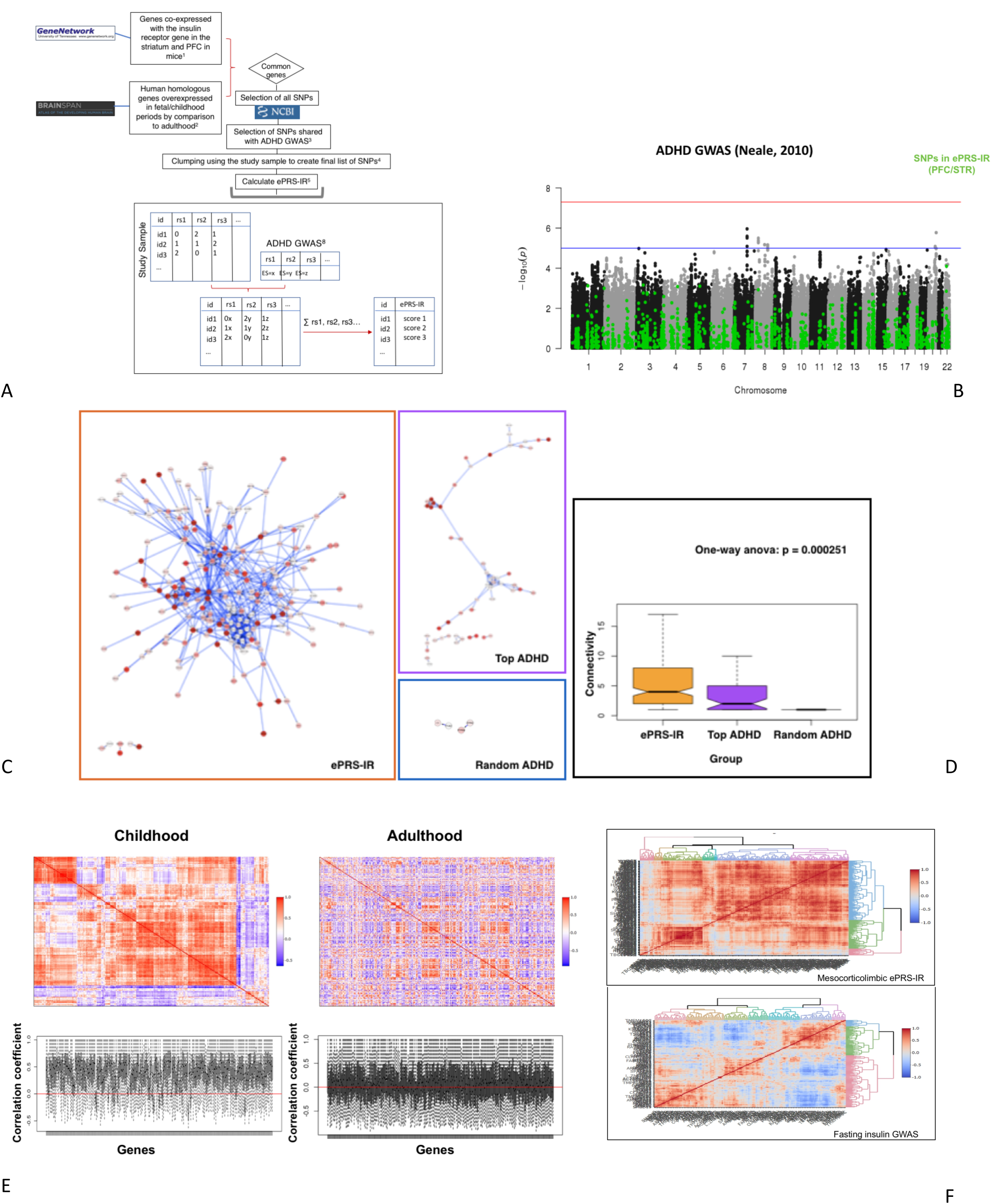
The mesocorticolimbic ePRS-IR. **A)** Flowchart depicting the steps involved in creating the expression-based and mesocorticolimbic-specific polygenic risk score based on genes co-expressed with the insulin receptor (ePRS-IR) using gene co-expression databases. i) GeneNetwork was used to generate a co-expression matrix with insulin receptor (IR) in the ventral striatum and in the prefrontal cortex in mice (absolute value of the co-expression correlation r≥0.5); ii) BrainSpan was then used to identify consensus autosomal transcripts from this list with developmental enrichment within the same brain areas, selecting transcripts differentially expressed at ≥1.5 fold during child and fetal development as compared to adult samples. The final list included 281 genes; iii) Based on their functional annotation in the National Center for Biotechnology Information, U.S. National Library of Medicine using GRCh37.p13, we gathered all the existing SNPs from these genes (total = 11,068) and subjected this list of SNPs to linkage disequilibrium clumping, to inform removal of highly correlated SNPs (r2>0.2) across 500kb regions, resulting in 1897 independent functional SNPs based on the children’s genotype data from MAVAN (Study Sample ids); iv) We used a count function of the number of alleles at a given SNP (rs1, rs2…) weighted by the effect size (ES) of the association between the individual SNP and ADHD. The sum of these values from the total number of SNPs provides the ePRS-IR score. References; 1. Mulligan, Mozhui et al. 2017, 2. Miller, Ding et al. 2014, 3. Neale, Medland et al. 2010, 4. Wray, Lee et al. 2014, 5. Chen, Yao et al. 2018. **B)** Single nucleotide polymorphisms (SNPs) included in the mesocorticolimbic ePRS-IR, shown in green, in relation to the Manhattan plot of the ADHD GWAS(Neale, Medland et al. 2010). The biologically-informed method for selecting SNPs is designed to capture signals associated with a functional gene network, and hence it can be expected that none of the individual SNPs included in the score are statistically significantly associated with the disease (in this case, ADHD). **C)** Gene Network analysis of the mesocorticolimbic ePRS-IR (larger panel), random (top) and ADHD 2010 (bottom small panel) 281 genes. The random gene list was created selecting a random group of SNPs of the same number of SNPs included in the ePRS-IR. The ADHD 2010 gene list was created selecting genes associated with the most significant SNPs from the ADHD GWAS (Neale, Medland et al. 2010). The picture demonstrates that the ePRS-IR represents a network with significantly higher connectivity than a list of genes associated with a random list of SNPs, and also in comparison to the top genes from the ADHD GWAS. **D)** Comparison of the number of connections between the genes in each network (mesocorticolimbic ePRS-IR, random list and top genes in the ADHD GWAS). Genes included in the mesocorticolimbic ePRS-IR have significantly more interactions than the random list, and the top genes from the ADHD GWAS (One-Way ANOVA, p<0.05). This suggests a more cohesive gene network of biological relevance. **E)** Coexpression of the genes included in the mesocorticolimbic ePRS-IR, in childhood and adulthood, combining gene expression data from PFC and striatal regions. Top panels: The heatmap of the genes’ co-expression in childhood (left) shows several clusters of highly co-expressed genes. Although the clusters are not maintained in the retained order heatmap for adulthood (right), some other clusters of co-expressed genes are observed. Bottom panels: values of the gene expression correlation coefficients in childhood (left) and adulthood (right). Each vertical line represents correlation with a unique gene. The red line is drawn at a correlation score of zero. Especially in childhood, most genes included in the score have highly correlated gene expression values. Data for this analyses were extracted from BrainSpan (Miller, Ding et al. 2014). **F)** Co-expression of the genes included in the mesocorticolimbic ePRS-IR (top panel) and a comparable number of genes associated with the most significant SNPs from the fasting insulin GWAS (Manning, Hivert et al. 2012) (bottom panel), from birth to 11 years of age in the PFC and striatal regions. The heatmaps demonstrate that while the mesocorticolimbic ePRS-IR includes genes that are highly co-expressed in striatum and PFC, genes from the fasting insulin GWAS are less consistently co-expressed in these brain regions. This suggests that the ePRS methodology captures gene networks that are cohesive in specific brain regions, which may not be represented even when analyzing a genetically correlated GWAS (fasting insulin).

This biologically-informed method for selecting SNPs is designed to capture risk associated with a functional unit of a gene network, and hence it can be expected that none of the individual SNPs included in the score are statistically significantly associated with ADHD diagnosis in the GWAS (Fig. 1B), but instead, the cumulative ePRS-IR score associates with the relevant endophenotypes. A gene network analysis was performed comparing the list of the 281 genes included in the ePRS-IR with (a) the genes associated with a random selection of 11,068 SNPs from the ADHD GWAS (Neale, Medland et al. 2010) and (b) the top 281 genes associated with the most significant SNPs from the ADHD GWAS (Neale, Medland et al. 2010) (Fig. 1C). The analysis shows that the number of interactions between the genes that comprise the ePRS-IR is significantly higher than the control random list, or the genes from the ADHD GWAS (Fig. 1D), suggesting that the biologically-informed score represents a much more cohesive gene network. We examined the co-expression patterns of the genes included in the mesocorticolimbic ePRS-IR over the life-course using data from BrainSpan (Miller, Ding et al. 2014). Several genes are highly co-expressed in childhood, as expected from the methodological approach used to select the genes included in the score. This co-expression pattern is not maintained into adulthood, although other clusters of co-expression emerge (Fig. 1E). As a validation of our mesocorticolimbic ePRS-IR score, we created 30 random ePRS and repeated linear regression analyses for each of the random ePRSs. The results of the regression analyses involving different ePRS scores were pooled together to obtain an estimate of the effect of the ePRS interaction with sex on the IST performance at 72 months. The results showed a non-significant interaction term (fixed condition: =0.38, p =0.81; decreasing condition =0.14, p =0.92). When analyzing the shared heritability between our ePRS-IR and other complex traits using a LD score regression (Bulik-Sullivan, Loh et al. 2015), there was a significant genetic correlation between the GWAS for cross-disorder (Cross-Disorder Group of the Psychiatric Genomics 2013) and the mesocorticolimbic ePRS-IR (rg=1.1, p=0.004), which was expected given that the ADHD GWAS is part of the cross-disorder genome-wide analysis. However, we observe the brain specificity of the mesocorticolimbic ePRS-IR in relation to peripheral insulin levels or sensitivity, as there was no significant shared heritability between the ePRS-IR and the fasting insulin GWAS(Manning, Hivert et al. 2012) (rg= 0.53, p=0.25) or HOMA-IR GWAS (Dupuis, Langenberg et al. 2010)(rg=0.62, p=0.16). Brain specificity is also demonstrated when comparing the co-expression patterns of the genes included in our score and an equivalent number of genes associated with the most significant SNPs from the fasting insulin GWAS (Scott, Lagou et al. 2012) in the PFC and striatal regions (Fig. 1F). While the mesocorticolimbic ePRS-IR has highly co-expressed genes in these brain areas, genes from the fasting pro-insulin GWAS are less consistently co-expressed in these regions. This confirms the ability of the ePRS methodology to generate a genetic score that denotes tissue-specific gene networks, and highlights the meaningful difference between the information contained in the ePRS-IR score and peripheral levels of metabolic hormones such as insulin.

### Endophenotypic differences predicted by the mesocorticolimbic ePRS-IR

We then analyzed whether the ePRS-IR would predict impulsivity, a highly sex-specific trait, in 6-year old children (O’Donnell 2014) tested using a computer-based, Information Sampling task (IST). There was a significant interaction effect between ePRS-IR and sex on the Information Sampling Task (IST) (fixed condition = 2.385, p = 0.016; decreasing condition = 2.479, p = 0.006) applied at 72 months (Fig. 2A and B). While a simple slopes analysis showed no relationship between the ePRS-IR score and mean P Correct values in girls (fixed condition = −0.636, p = 0.36; decreasing condition = −0.91, p = 0.15), a lower ePRS-IR was significantly related to lower mean P Correct (less certainty when coming up to a decision or higher reflexive impulsivity) in boys (fixed condition = 1.75, p = 0.01; decreasing condition = 1.57, p = 0.01). We next examined whether a “random” PRS or other conventional PRSs (for ADHD (Neale, Medland et al. 2010) or addiction (Tobacco and Genetics 2010)) of comparable size in terms of SNP number could also predict impulsivity in these children. None of these control scores was associated with the interaction effect on reflection impulsivity in this sample (Fig. 2C). We then hypothesized that a lower ePRS-IR, associated with childhood impulsivity in boys as shown above, would predict risk for addiction in men, considering the clinical overlap between impulsive phenotypes and risk for addiction (Fatseas, Hurmic et al. 2016). We therefore investigated if the ePRS-IR was associated with the risk for addiction phenotypes using data from the Study of Addiction: Genetics and Environment (SAGE) repository(Bierut, Saccone et al. 2002). The analysis revealed a significant interaction between the ePRS-IR score and sex for the number of addiction comorbidities (interaction effect; = −11.33, p = 0.01) and probability of alcohol dependence (interaction effect; = −13.71, p = 0.02). Further simple slopes analysis confirmed that a lower ePRS-IR was significantly associated with number of addiction comorbidities only in males (males, simple slope = −7.30, p = 0.04; females, simple slope = 4.59, p = 0.14); simple slopes analysis for alcohol dependence was marginally significant in males (= −7.99, p = 0.06, OR=3.38) with no effect in females (= 6.40, p = 0.10) (Fig. 2D and E) (Table S2).

**Fig. 2.**
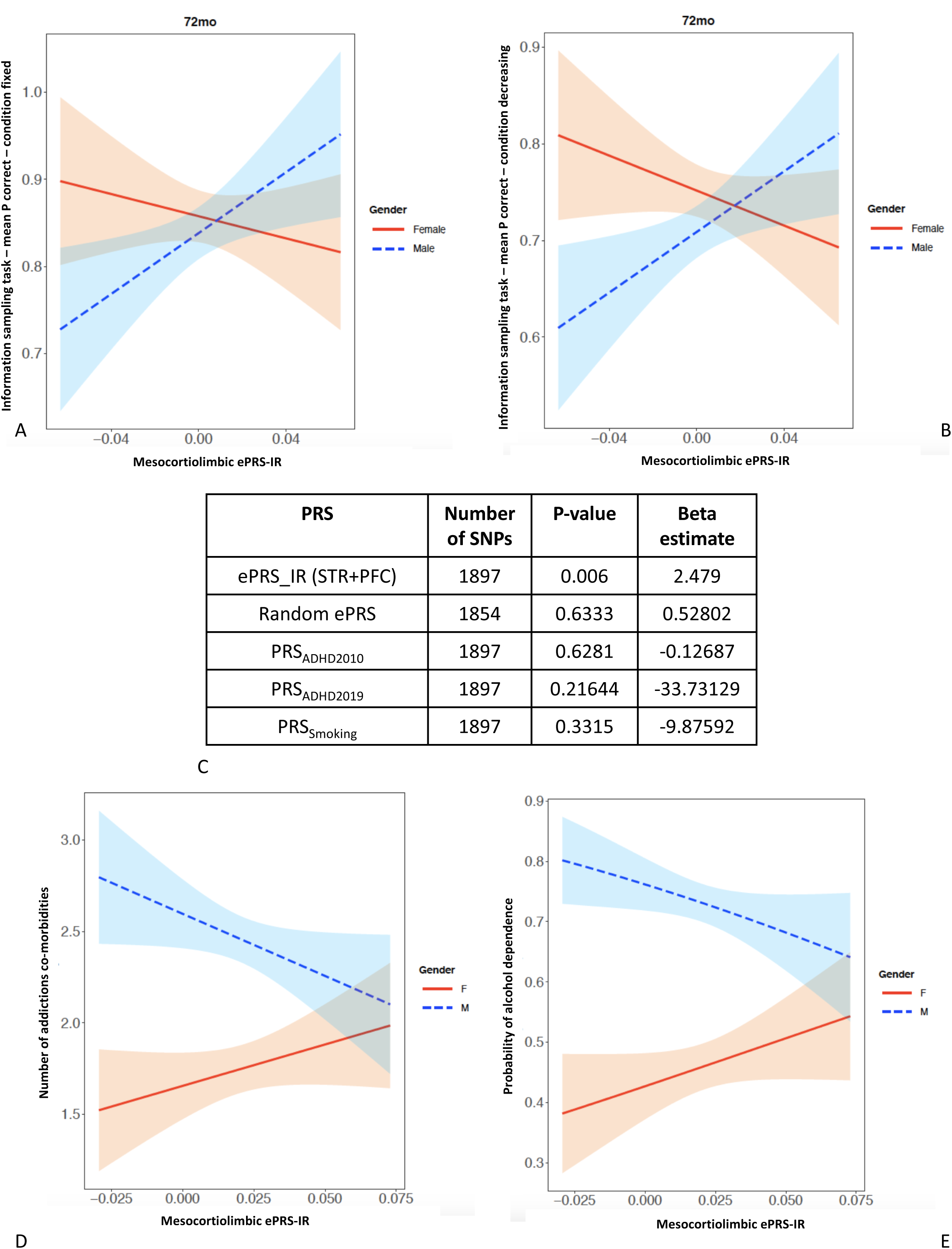
Phenotypic differences predicted by the mesocorticolombic ePRS-IR. **A) and B)** Performance in the Information Sampling Task (IST, CANTAB) at 72 months according to sex and ePRS-IR in fixed (A) and decreasing (B) conditions. There is a significant interaction between the genetic score and sex on IST outcome, in which boys with lower ePRS-IR sample less information before taking the decision, being more impulsive than boys with higher ePRS-IR. Boys are depicted in blue and girls in red. N=207. **C)** Validation of the findings by using different polygenic risk scores to investigate the interaction with sex on the Information Sampling Task in children. The mesocorticolimbic ePRS-IR significantly interacted with sex as shown in Figure 2A, predicting impulsivity in boys. A PRS with a random SNP selection, a classic PRS from two different GWASes for ADHD (Neale, Medland et al. 2010, Demontis, Walters et al. 2019) or a PRS for addiction (i.e. smoking) (Tobacco and Genetics 2010) were unable to predict the behavioral phenotype. **D)** Number of addiction co-morbidities according to sex and mesocorticolimbic ePRS-IR in the SAGE cohort. There is a significant interaction between the genetic score and sex on the number of co-morbidities, in which ePRS-IR has a negative effect on the number of co-morbidities in men but not women. **E)** Probability of alcohol abuse in the SAGE cohort. A significant interaction between the genetic score and sex was found on the probability of alcohol dependence, in which ePRS-IR has a negative effect on the probability of alcohol dependence in men but not women.

### Comparison between mesocorticolimbic ePRS-IR using two versions of the ADHD GWAS - 2010 (Neale, Medland et al. 2010) and 2019 (Demontis, Walters et al. 2019)

Considering the theoretical premise of the ePRS-IR (biological polygenic score based on gene co-expression), we hypothesized that the choice of the GWAS for providing effect sizes for weighing the SNPs would not have a major influence on the prediction ability of the expression based polygenic score. We then created simultaneously two mesocorticolimbic ePRS-IR scores at the point of the SNP selection, weighing one using the 2010 ADHD GWAS (Neale, Medland et al. 2010) and the second one using the recently published 2019 ADHD GWAS (Demontis, Walters et al. 2019). Confirming our hypothesis, both scores were associated with a significant interaction effect with sex on impulsivity phenotype measured by the IST task in MAVAN (2010: = 1.13, p = 0.05; 2018: = 128.8, p = 0.05) (Figure S1). Considering that several studies described associations between polygenic scores using the 2010 ADHD GWAS and endophenotypes associated with attention-impulsivity problems in community samples (rather than ADHD cases vs. controls, similar to our main cohort MAVAN)(Groen-Blokhuis, Middeldorp et al. 2014, Martin, Hamshere et al. 2014), we opted for our main analysis to be focused on the 2010 ADHD GWAS.

### The hippocampal ePRS-IR

To further validate the brain-specificity of the biologically-informed polygenic score, we followed the same approach depicted in Fig 1A and created the hippocampal ePRS-IR by changing the brain region of interest in the process, and compiling the SNPs using the association betas described in the Alzheimer’s GWAS (Lambert, Ibrahim-Verbaas et al. 2013) (Table S3). Similar to the mesocorticolimbic ePRS-IR, the selected SNPs are not statistically significantly associated with Alzheimer’s in the GWAS (Fig. 3A). The gene network analysis demonstrates that the hippocampal ePRS-IR also represents a highly interconnected network (Fig. 3B). Analysis of the co-expression pattern of the genes included in the hippocampal ePRS-IR revealed that the high co-expression clusters from childhood are not maintained in adult life (Fig. 3C). Informed by the role of the IR in the hippocampus on cognition, we explored this endophenotype in children by evaluating cognitive performance on the Number Knowledge Task applied at 48 months (O’Donnell 2014). There was a main effect of the hippocampal ePRS-IR in predicting task performance (p = 0.03, = −1350.1), in which children with higher ePRS-IR scores have lower performance on the cognitive test (Fig. 3D). We found no significant interaction effect between the hippocampal ePRS-IR and sex on the task at this age. The hippocampal ePRS-IR was also able to distinguish the presence of Alzheimer’s disease diagnosis in the GenADA study (Li, Wetten et al. 2008) (p = 6.82e-09, = 972.40), in which there are more cases in individuals with higher ePRS-IR scores (Fig. 3E). No significant interaction between ePRS and sex was found in the adult cohort.

**Fig. 3.**
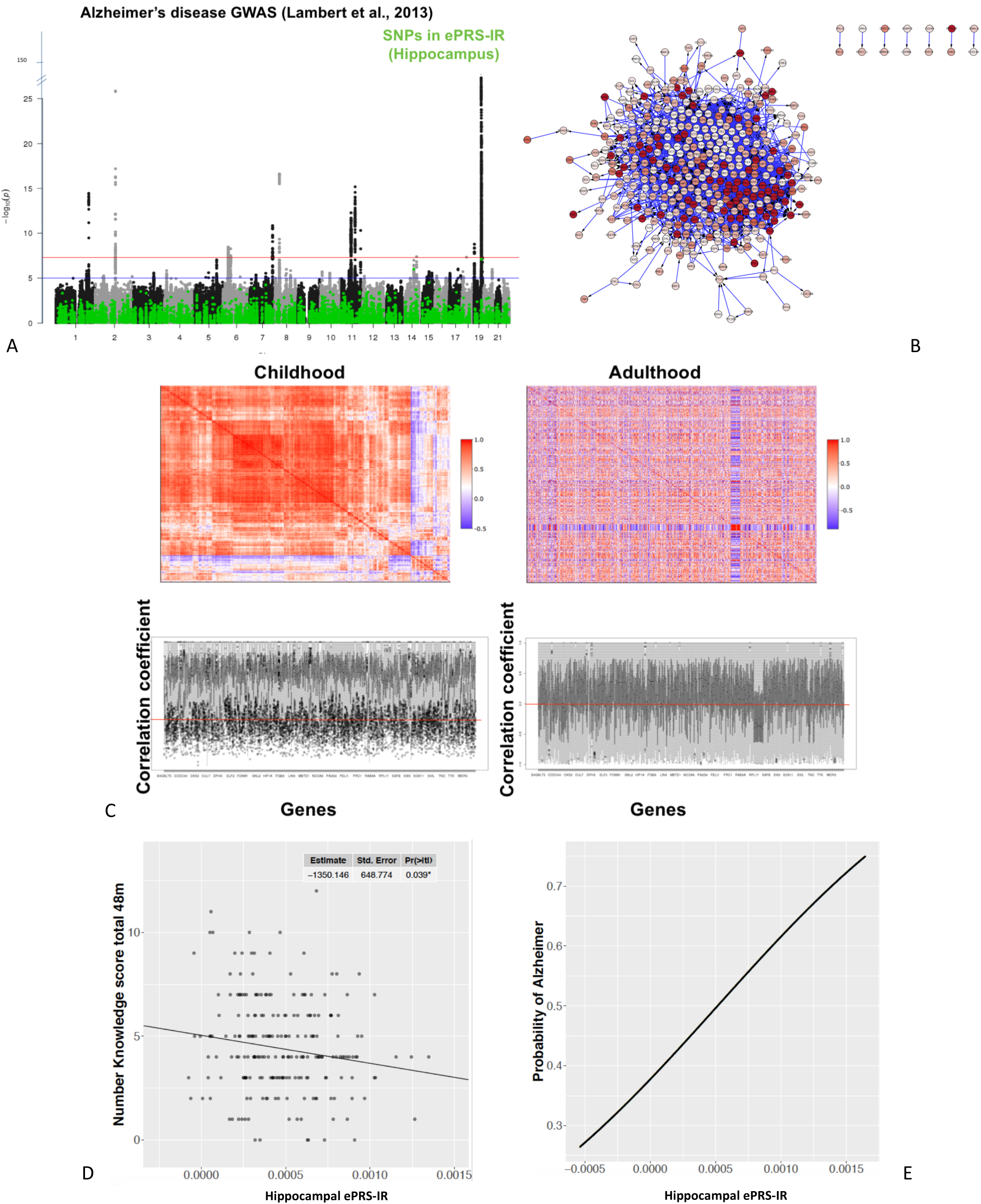
The hippocampal ePRS-IR. **A)** Single nucleotide polymorphisms (SNPs) included in the hippocampal ePRS_IR, shown in green, in relation to the Manhattan plot of the Alzheimer’s GWAS (Lambert, Ibrahim-Verbaas et al. 2013). The SNPs captured in the score do not have genome-wide significance. **B)** Gene network analysis in the hippocampal ePRS-IR 543 genes. The picture demonstrates that the hippocampal ePRS-IR also represents a highly cohesive gene network. **C)** Coexpression of the genes included in the hippocampal ePRS-IR, in childhood and adulthood. Top panels: The heatmap of the genes’ co-expression in childhood (left) shows several clusters of highly co-expressed genes. The clusters are not maintained in the retained order heatmap for adulthood (right) suggesting that these gene networks are relatedly connected in childhood, but not anymore in adulthood. Bottom panels: values of the correlation coefficients of the gene expression values in childhood (left) and adulthood (right). Each vertical line represents correlation with a unique gene. The red line is drawn at a correlation score of zero. Genes included in the score have highly correlated gene expression in childhood, but this is not seen in adult samples. **D)** Main effect of hippocampal ePRS-IR on Number Knowledge Task performance, applied at 48 months, in which children with higher ePRS-IR scores have lower performance on the cognitive test. No significant interaction between ePRS and sex was found at this age. **E)** Presence of Alzheimer’s disease diagnosis according to hippocampal ePRS-IR scores in the GenADA study. There was a main effect of the hippocampal ePRS-IR in predicting Alzheimer’s cases in which there are more cases in individuals with higher ePRS-IR scores. N=1565.

### Comparison between mesocorticolimbic and hippocampal ePRS-IR scores

To examine the isolated contribution of unique SNPs to the above described effects, we performed a SNP by sex association analysis for the mesocorticolimbic ePRS-IR (investigating the interaction with sex) and a genome wide association analysis for the hippocampal ePRS-IR (investigating the main effects of ePRS-IR) using linear regressions. There was no unique SNP included in the mesocorticolimbic ePRS-IR responsible for the SNP by sex interaction on impulsivity in children, or early addiction onset in adults. Similarly, no isolated SNP included in the hippocampal ePRS-IR was associated with poorer cognitive performance in children, or Alzheimer’s disease, confirming that variations in the biological function of the gene networks represented in the ePRS-IR scores predict their respective outcomes as a global score, with small contributions from all included SNPs, but not by any isolated mutations (Fig. 4A). To further validate the brain region specificity of the scores, we analyzed the distribution of children in each quartile of the mesocorticolimbic and hippocampal ePRS-IR in MAVAN (Fig. 4B) and verified that the scores are independent (r=0.062, p=0.416). In addition, the mesocorticolimbic ePRS-IR did not predict cognitive performance (p = 0.08, = −12.36 for main effect, p = 0.62, = −6.93 for the interaction with sex). Likewise, the hippocampal ePRS-IR was not associated with IST scores (p = 0.11, = 0.02 for main effect, p = 0.29, = −0.020 for the interaction with sex). Enrichment analysis of the SNPs comprising the mesocorticolimbic ePRS-IR using Metacore^®^ (Thomson Reuters) showed statistically significant enrichment for pathways involved in cell cycle regulation (FDR q = 1.18e-02). Process networks were significant for neurogenesis/axonal guidance (FDR q = 6.667e-03), transcription and translation (FDR q = 1.39e-02). The hippocampal ePRS-IR is enriched for pathways involved in development (hedgehog signaling, FDR q=1.19e-05) and transcription (FDR q=1.42e-04). Process networks involve cell division (Fig. 4C). Both ePRS-IRs were significantly associated with similar transcription factors including CREB1, C-Myc, SP1 and ESF1 (Fig. 4D).

**Fig. 4.**
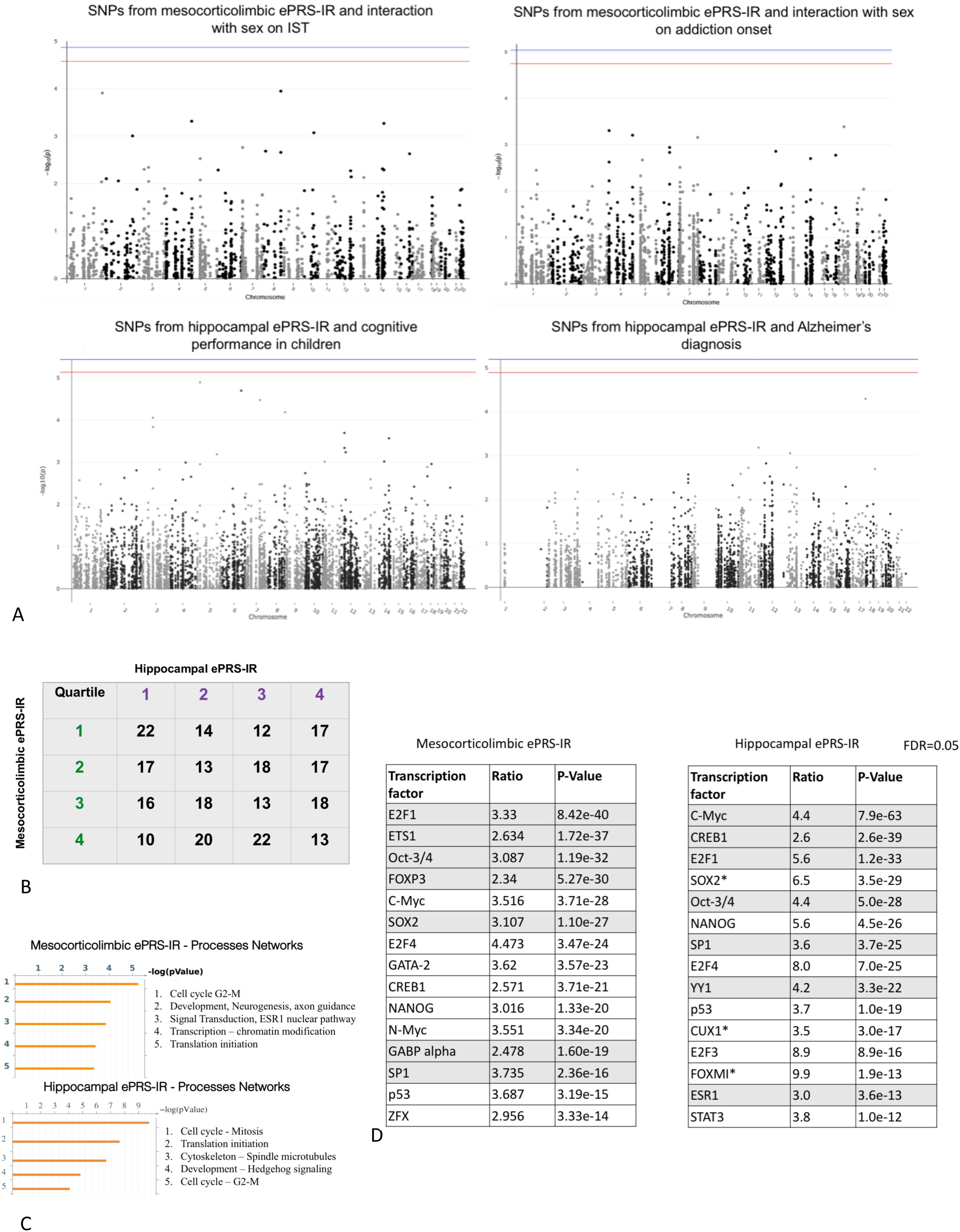
ePRS-IR scores are brain-region specific. **A)** Manhattan plots investigating the contribution of isolated SNPs into the reported findings. There was no unique SNP included in the mesocorticolimbic ePRS-IR responsible for the SNP by sex interaction on impulsivity in children (top left) or alcohol dependence in adults (top right). Similarly, no isolated SNP included in the hippocampal ePRS-IR was associated with poorer cognitive performance in children (bottom left), or Alzheimer’s disease (bottom right). These findings confirm that variations in the biological function of the gene networks represented in the mesocorticolimbic ePRS-IR and hippocampal ePRS-IR can predict these outcomes as a global score, with small contributions from all included SNPs, but not by any isolated mutations. **B)** Pearson’s product-moment correlation demonstrating the distribution of children in each quartile of the mesocorticolimbic and hippocampal ePRS-IRs. There is no correlation between the quartiles of the two ePRSs (r=0.062, p=0.416). This shows that a subject with a high mescorticolimbic ePRS-IR would not necessarily have a high score in the hippocampal ePRS-IR, and vice-versa, demonstrating their brain specificity and independence. **C)** Enrichment analysis of mesocorticolimbic (upper panel) and hippocampal (bottom panel) ePRS-IR. **D)** Transcription factor analysis of the two brain-region specific ePRS-IRs.

## Discussion

There has been a marked increase in the number of GWAS available. We now have access to association studies addressing a range of health related-disease and otherwise-outcomes. Conventional polygenic risk scores derived from existing GWAS’s are intended to reflect genetic vulnerability. However, the PRS approach is limited by the small explained variance in GWAS of common disorders and a lack of insight into the phenotypic constructs of the disease. We propose an alternative approach that integrates the power of GWAS datasets to form an ePRS informed by research in neuroscience, as well as the strength of transcriptomic datasets. The resulting brain-region specific ePRS reflects an integrated gene network and molecular pathway that allows the use of genotyping data to test candidate biological mechanisms (Fig. 5).

**Fig. 5.**
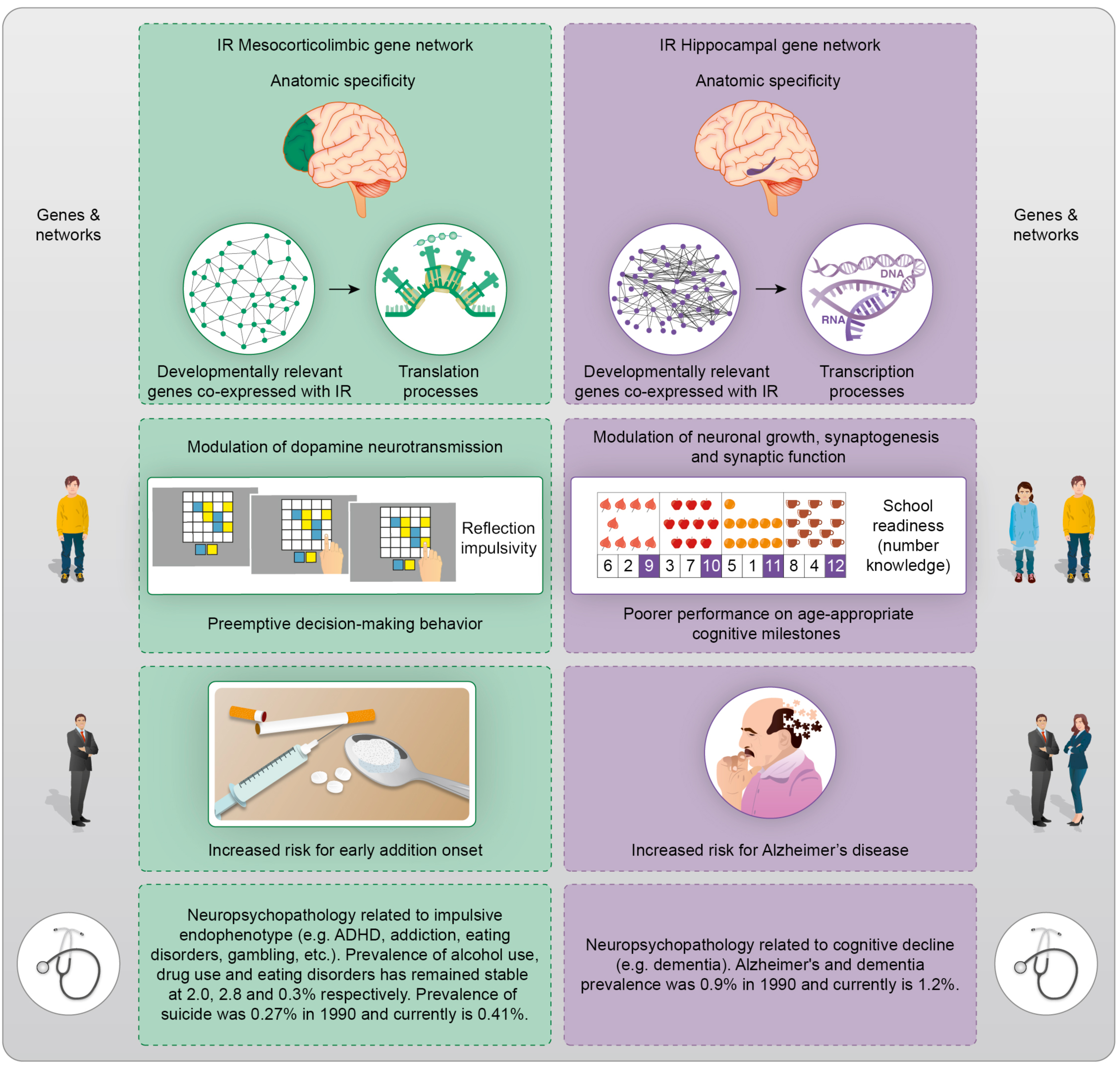
Theoretical framework and applications. The expression based, brain region specific ePRS-IR represents a developmentally relevant gene network with enrichment for specific processes like translation and transcription. Variation in this score predicts particular endophenotypes in childhood – for example, impulsivity and cognitive performance. These behaviors map onto risk for diseases later in life, such as early addiction onset and Alzheimer’s disease, and the eRPS-IR is equally associated with these outcomes. Psychopathology related to poor inhibitory control or decision-making encompass a wide range of conditions (ADHD, addiction, eating disorders, gambling, suicide, etc.). Similarly, there is an increasing incidence of Alzheimer’s disease and other dementias. The neurobiological understanding of these conditions contributes to the development of tools for early identification and primary prevention.

The conventional polygenic profiling method is driven by a statistical approach. Additional levels of biological complexity, such as biomarkers and neuroanatomic loci of relevance - from experimental or theoretical databases - can be superimposed and inform the use of this technology, as we demonstrate here. The need to consider gene networks is further exemplified by the observation that the power to detect a causal SNP is drastically reduced when the trait/disorder is caused by two or more pathways (Wray and Maier 2014). We showed that a biologically-informed polygenic risk score based on genes co-expressed with the central IR represents highly cohesive and relevant gene networks. The mesocorticolimbic-specific ePRS-IR is more strongly associated with impulsivity and the risk for early onset of substance dependence in males than is a conventional PRS for either ADHD or addiction. The sex-specificity of our findings was expected, given the increased prevalence of ADHD and behavioral alterations associated with this condition (such as impulsivity) in boys compared to girls (Silveira, Agranonik et al. 2012, Vasiliadis, Diallo et al. 2017). Likewise the hippocampal-specific ePRS-IR was associated both with cognitive performance in children and with Alzheimer’s diagnosis in later ages, demonstrating that the biologically-informed framework is brain region and endophenotype-specific.

Our network analysis shows that the ePRS-IR represents a cohesive gene network with significantly more connections than the list of genes extracted from the GWAS for ADHD, or random lists SNPs. This robust approach therefore goes beyond describing associations between single gene variants and the outcomes, but captures information about the whole gene network, and its function, in specific brain regions. One of the critiques of GWAS is that they are ‘a static global’ measure, unable to discern spatial or temporal differences, as compared to techniques such as RNAseq or WGBS, which are fundamental to the study of any trait/disorder. Our approach integrates a static global measure-which is easy to acquire in your dataset of interest-with a temporal and spatial information -which are, usually, not feasible to acquire in all human datasets - to compute a score that is adept at informing spatially relevant endophenotypes.

Conduct disorder and impulsivity are the foremost risk traits for alcohol use disorder among the 80 personality disorder criteria of DSM-IV (Rosenstrom, Torvik et al. 2017). There is a relationship between childhood ADHD and the risk for developing drug addiction later in life (Levy, Katusic et al. 2014), especially considering the impulsivity component, rather than inattention (Sihvola, Rose et al. 2011), in agreement to the findings described here. Insulin function is associated with the risk for drug addiction (O’Dell and Nazarian 2016). Diminished insulin sensitivity is related to less endogenous dopamine at D2/3 receptors in the ventral striatum(Caravaggio, Borlido et al. 2015), reinforcing the idea that metabolic processes are involved in dysfunctions of the mesocorticolimbic system, such as drug dependence. The enrichment analysis of our mesocorticolimbic ePRS-IR is consistent with the known role of insulin in cell division (Perdereau, Cailliau et al. 2015), neurogenesis (Ronaghi, Zibaii et al. 2019), signal transduction (Pearson-Leary, Jahagirdar et al. 2018), and axon guidance (Gupta, Yadav et al. 2018).

The hippocampal formation is the region with the higher IR gene expression in the brain (De Felice and Benedict 2015). Lower expression of IR and altered levels of different components of the insulin signaling pathways were described in hippocampi from Alzheimer’s patients (Talbot, Wang et al. 2012). A down-regulation in insulin signaling causes memory impairment through inhibition of serine phosphorylation of the insulin receptor via activation of stress kinases like c-Jun-n-terminal kinase (JNK)(Bomfim, Forny-Germano et al. 2012). JNK enhances glycogen synthase kinase 3β (GSK3β) activity that leads to tau phosphorylation followed by accumulation of neurofibrillary tangles – a hallmark of Alzheimer’s disease - in the brain (Bedse, Di Domenico et al. 2015). Others have shown that impaired insulin sensitivity is linked to cognitive deficits in the elderly (Benedict, Brooks et al. 2012), and that measures of cognitive function in youth predict later risk for Alzheimer’s disease (Snowdon, Greiner et al. 2000, Bloss, Delis et al. 2008), in alignment to our current results.

Our genomic approach integrates information from molecular neurobiology with GWAS technology to develop a biologically-informed polygenic score based on gene co-expression data from specific brain regions. This approach creates a novel genomic measure to identify genetic vulnerability for childhood behavioral phenotypes that predict later neuropsychiatric conditions in community-based samples, highlighting possible targets for drug development.#add a closing clinical statement here#.

## Author contributions

Conceptualization PPS, MJM, SHD; data curation KM, IP, LMC, TTTN, EG, LMM, JLM; formal analysis IP, SHD, PPS; funding acquisition MJM, MK; investigation PPS, KM, SHD; methodology PPS, SHD, KOD, MY; project administration JD; resources LMM, JLM, EG, TTTN; supervision MJM, PPS; validation SHD, LMC, MY; visualization SHD, KM, LMC; writing – original draft PPS, SHD, MJM; writing – review & editing all authors;

## Data and materials availability

Data and materials are available upon request.

## Supplementary Materials

Materials and Methods

Figure S1

Tables S1 to S3

