## Supplementary material for "A biologically-informed polygenic score identifies endophenotypes and clinical conditions associated with the insulin receptor function on specific brain regions"

A biologically-informed polygenic score of the insulin receptor gene network

**This PDF file includes:**

Materials and Methods

Tables S1 to S3

Figure 1

Materials and Methods

Samples

Main Cohort: We used data from the prospective Maternal Adversity, Vulnerability and Neurodevelopment (MAVAN) birth cohort (O'Donnell 2014) that followed children at different time points in the first years of life in Montreal (Quebec) and Hamilton (Ontario), Canada. Approval for the MAVAN project was obtained from obstetricians performing deliveries at the study hospitals and by the ethics committees and university affiliates (McGill University and Université de Montréal, the Royal Victoria Hospital, Jewish General Hospital, Centre hospitalier de l’Université de Montréal, Hôpital Maisonneuve-Rosemount, St Joseph’s Hospital and McMaster University). Informed consent was obtained from all subjects.

Replication Cohorts: A) Study of Addiction, Genetics and Environment (SAGE) repository(Bierut, Saccone et al. 2002, Edenberg 2002, Edenberg, Xuei et al. 2006, Bierut 2007, Bierut, Madden et al. 2007, Saccone, Hinrichs et al. 2007, Bierut, Strickland et al. 2008), acquired from dbGaP (https://www.ncbi.nlm.nih.gov/gap, Accession number: phs000092.v1.p). The SAGE dataset was compiled from three studies: the Collaborative Study on the Genetics of Alcoholism (COGA); the Family Study of Cocaine Dependence (FSCD); and the Collaborative Genetic Study of Nicotine Dependence (COGEND). The SAGE dataset contains genotyping and clinical phenotypes related to substance dependence for adult subjects. We received access to the SAGE dataset based on the approval of our Data Access Request (DAR) by the NIH Data Access Committee. We agree with the stipulations of the Data Use Certification. B) GEN-ADA(Li, Wetten et al. 2008, Filippini, Rao et al. 2009) is a multi-site study conducted at GlaxoSmithKline Inc and nine medical centres. The study was designed to capture genetic information from 1000 Alzhiemiers’ disease patients and 1000 matched controls. We received access to the GenADA dataset based on the approval of our Data Access Request (DAR) by the NIH Data Access Committee. We agree with the stipulations of the Data Use Certification.

Genotyping:

In MAVAN, we genotyped 242,211 autosomal SNPs using genome-wide platforms (PsychArray/PsychChip, Illumina) according to manufacturer’s guidelines with 200ng of genomic DNA derived from buccal epithelial cells and our quality control procedures. Specifically, we removed SNPs with a low call rate (<95%), low p-values on Hardy-Weinberg Equilibrium exact test (p<1e-40), and minor allele frequency (<5%). Afterward, we performed imputation using the Sanger Imputation Service (McCarthy, Das et al. 2016) resulting in 20,790,893 SNPs with an info score >0.80 and posterior genotype probabilities >0.90. In SAGE, we used the imputed genotypes provided in the data repository. The imputation was performed using BEAGLE. Post imputation, SNPs were filtered to only include those with an r^2^>0.3 (total = 4,566,998). In GenADA, genotyping data was provided for 402,708 SNPs, we removed SNPs with a low call rate (<95%), low p-values on Hardy-Weinberg Equilibrium exact test (p<1e-40), and minor allele frequency (<5%). Afterwards, we performed imputation using the Sanger Imputation Service (McCarthy, Das et al. 2016), resulting in 7,267,018 SNPs with an info score >0.80 and posterior genotype probabilities >0.90.

ePRS-IR calculation:

The polygenic risk score based on genes co-expressed with the insulin receptor (ePRS-IR) was created using gene co-expression databases including 1) GeneNetwork ([http://genenetwork.org](http://genenetwork.org/)), 2) BrainSpan (http://brainspan.org), 3) NCBI Variation Viewer (https://www.ncbi.nlm.nih.gov/variation/view/). These resources allowed us to identify genes co-expressed with the IR in the striatum and prefrontal cortex (PFC) regions in mice (GeneNetwork) and humans (BrainSpan), and to identify SNPs for these genes in humans (NCBI Variation Viewer). The PRS was constructed as follows: 1) we used GeneNetwork to generate co-expression matrix with IR in the i) ventrial striatum, ii) PFC in mice (absolute value of the co-expression correlation r≥0.5), 2) we then used BrainSpan to identify consensus transcripts from this list with a child and fetal enrichment within the human brain. We selected autosomal transcripts differentially expressed in these brain regions at ≥1.5 fold during child and fetal development as compared to adult samples(Miller, Ding et al. 2014). The final list included 281 genes. Based on their functional annotation in the National Center for Biotechnology Information, U.S. National Library of Medicine (https://www.ncbi.nlm.nih.gov/variation/view/) using GRCh37.p13 we gathered all the existing SNPs from these genes (total = 11,068) and subjected this list of SNPs to linkage disequilibrium clumping, which uses the lowest association p-values in the ADHD GWAS to inform removal of highly correlated SNPs (r^2^>0.2) across 500kb regions(Wray, Lee et al. 2014), resulting in 1897 independent functional SNPs based on the children’s genotype data from MAVAN. We compared the distributions of the score in the two cohorts (MAVN and SAGE) using Kolmogorov-Smirnov test and there were no significant differences between the scores (p=0.996). We used a count function of the number of alleles at a given SNP weighted by the effect size of the association between the individual SNP and ADHD(Neale, Medland et al. 2010). All SNPs were subjected to linkage disequilibrium clumping (r^2^>0.2 across 500kb) so only independent SNPs that are most associated to ADHD, based on the association p-values in the ADHD GWAS, comprised the PRS.

The hippocampal ePRS-IR was build using the same pipeline described above, having the hippocampus as the brain structure of interest. The final list included 544 genes (530 excluding genes on X and Y chromosomes) and 363,412 SNPs; this list was subjected to linkage disequilibrium clumping, resulting in 6,594 dependent functional SNPs based on the children’s genotype data from MAVAN. We compared the distributions of the score in the two cohorts (MAVAN and GenADA) using Kolmogorov-Smirnov test and there were no significant differences between the scores (p=0.668). We used a count function of the number of alleles at a given SNP weighted by the effect size of the association between the individual SNP and Alzheimer’s disease (Lambert, Ibrahim-Verbaas et al. 2013).

Behavioral phenotyping

Reflexive impulsivity in children: The Information Sampling Task (IST) from the CANTAB was designed to measure reflection impulsivity and decision making(Luciana and Nelson 2002), and was applied at 72 months in MAVAN. Rather than relying on speed-accuracy indices, IST measures reflection impulsivity by calculating the probability of the subject selecting the correct answer after making a decision based on the information sampled prior to making that decision. On each trial, children are presented with a 5×5 matrix of grey boxes on the computer screen, and two larger colored panels at the foot of the screen. They are told that it is a game for points, won by correctly choosing the color under the majority of the grey boxes. Touching a grey box immediately opens that box to reveal one of the two colors displayed at the bottom of the screen. Subjects could open boxes at their own will with no time limit before deciding between one of the two colors, indicating their decision by touching one of the two panels at the bottom of the screen. When they do, the remaining boxes are uncovered and a message is displayed to inform them whether or not they were correct. The primary performance outcome measure was the mean probability of being correct at the point of decision (P Correct). P Correct is the probability that the color chosen by the subject at the point of decision would be correct, based only on the evidence available to the subject at the time (i.e., dependent on the amount of information they had sampled). There was a recent update in the mean P Correct formula, which was endorsed by the original authors of the measure(Bennett, Oldham et al. 2016, Clark and Robbins 2016) , therefore in this study we calculated and used the new mean P Correct (Pokhvisneva, Léger et al. 2018).

Number Knowledge Task (NK): The Number Knowledge task is part of the School Readiness Battery, applied at 48 months in MAVAN, assessing school readiness which may be defined as the minimum developmental level allowing the child to respond adequately to school demands (Lemelin, Boivin et al. 2007). This task is a well validated diagnostic screening test of school readiness (Okamoto and Case 1996) (Okamoto & Case, 1996).

Addiction risk: The clinical assessment of substance dependency was based on a Semi-Structured Assessment for the Genetics of Alcoholism (SSAGA II)(Bucholz, Cadoret et al. 1994) and adapted versions of the SSAGAII, which assesses the physical, psychological and social manifestations of substance dependence. We used the number of addiction co-morbidities and the presence of alcohol dependence as outcomes.

Alzheimer’s diagnosis: The Alzheimer’s disease status was diagnosed using the Diagnostic and Statistical manual of Mental Disorders Fourth Edition (AmericanPsychiatricAssociation 2000)

Gene Network Analysis

We extracted a list of the 281 genes from the SNPs with the lowest p-values based on the post clumped results of the ADHD GWAS(Neale, Medland et al. 2010). RNA-sequencing data was downloaded from BrainSpan, including samples from 8 postconceptional weeks to 11 years old within prefrontal cortex (dorsolateral, ventrolateral, anterior cingulate cortex and orbitofrontal cortex) and striatum(Miller, Ding et al. 2014), for three gene lists: a) ePRS-IR; b) Random gene list (as detailed above) and c) ADHD top 281 genes. A median expression value was computed across the mentioned brain regions. The protein-protein interaction data were retrieved from STRING(Franceschini, Szklarczyk et al. 2013) (<https://string-db.org/)> and GeneMANIA(Warde-Farley, Donaldson et al. 2010) (<https://genemania.org>) databases and the protein-protein interaction networks were constructed and visualized in the Cytoscape software(Saito, Smoot et al. 2012). One-way ANOVA was used to compare these values across the three gene lists. The gene network for the hippocampal ePRS-IR was build using the same methodology. Enrichment and transcription factor analysis were performed using MetaCore™ (Clarivate Analytics).

Coexpression Analysis

We used publicly available gene expression data from BrainSpan(Miller, Ding et al. 2014) (http://www.brainspan.org) to analyze the correlation between the expression levels of the genes included in the ePRS-IR in the human mesocorticolimbic system and hippocampus, comparing children (1 to 12 years; n=26) to adults (20 to 40 years; n=22). The analyses were carried out in R using the heatmaply package.

Shared heritability

We performed a LD score regression (Bulik-Sullivan, Loh et al. 2015) regressing summary results statistics from the genetic variants across the genome on a measure of each variant’s ability to tag other variants locally, in order to estimate the genetic correlation between our mesocorticolimbic ePRS-IR and different complex traits and diseases using their GWAS summary-level results data (Bulik-Sullivan, Finucane et al. 2015), available in LD hub(Zheng, Erzurumluoglu et al. 2017).

Other genetic scores – analysis of validation

We generated other polygenic scores using our accelerated pipeline (<https://github.com/MeaneyLab/PRSoS>)(Chen, Yao et al. 2018), for each subject: (a) Random lists of SNPs from the ADHD GWAS (Neale, Medland et al. 2010); (b) PRSs for ADHD(Neale, Medland et al. 2010) (2010), ADHD (2019)(Demontis, Walters et al. 2019) and onset of smoking(Tobacco and Genetics 2010) of MAVAN children were computed. The PRSs are cumulative summary scores computed as the sum of the allele count weighted by the effect size across SNPs at a certain p-value thresholds (P_T_) based on the relevant GWAS(Wray, Lee et al. 2014). In all the genetic scores, the number of SNPs is comparable to the number of SNPs included in the mesocorticolimbic ePRS-IR.

Statistical Analysis:

Descriptive Statistics for all cohorts:

Statistical analysis of the baseline characteristics was performed using Spearman’s correlations, One-Way ANOVA ad Student’s T Test. The population structure was evaluated using principal component analysis of all autosomal SNPs that passed the quality control, without low allele frequency (MAF>5%) and are not in high linkage disequilibrium (r^2^>0.2) within a window of 50 SNPs at each step size of 5(Price, Patterson et al. 2006). Based on the inspection of the screeplot, the first three principal components were the most informative of population structure in this cohort and were included in all analyses.

Mesocorticolimbic ePRS-IR:

Linear regression analysis was performed to explore if the sex by ePRS-IR interaction was associated with the IST outcomes, adjusting for population structure and birth weight in MAVAN (Silveira, Pokhvisneva et al. 2018). Similarly, in SAGE, linear regression analysis was performed to explore if the sex by ePRS interaction was associated with the number of addiction co-morbidities or the presence of alcohol dependence (logistic regression in this case), adjusting for the population structure and study source. Simple slopes were analyzed to test the significance of the association in males and females when appropriate.

Hippocampal ePRS-IR:

In MAVAN, we explored the association between the hippocampal ePRS-IR and number knowledge task performance at 48 months of age using linear regression adjusted by population stratification and sex. In ADA, we used logistic regression to investigate the association between the hippocampal ePRS-IR and the presence of Alzheimer’s disease.

Data were analyzed using the Statistical Package for the Social Sciences (SPSS) version 20.0 software (SPSS Inc., Chicago, IL, USA) and R. We run all the analysis in the full datasets and repeated it excluding influential observations. As the results were similar, we chose to present the more conservative statistical approach excluding influential observations. Significance levels for all measures were set at α = 0.05.

Table S1.

List of genes selected for composing the mesocorticolimbic ePRS-IR (see details in the text).

| ***Gene Symbol*** | **Description** |
| --- | --- |
| *ACTL6A* | actin like 6A |
| *ACVR2B* | activin A receptor type 2B |
| *ADAMTS9* | ADAM metallopeptidase with thrombospondin type 1 motif 9 |
| *AGRN* | agrin |
| *AKT1* | AKT serine/threonine kinase 1 |
| *ARRDC3* | arrestin domain containing 3 |
| *ASF1B* | anti-silencing function 1B histone chaperone |
| *ASPM* | abnormal spindle microtubule assembly |
| *ATF7IP* | activating transcription factor 7 interacting protein |
| *BACH2* | BTB domain and CNC homolog 2 |
| *BCL11A* | B cell CLL/lymphoma 11A |
| *BCL11B* | B cell CLL/lymphoma 11B |
| *BCL9* | B cell CLL/lymphoma 9 |
| *BLMH* | bleomycin hydrolase |
| *BPHL* | biphenyl hydrolase like |
| *BRMS1* | breast cancer metastasis suppressor 1 |
| *CARHSP1* | calcium regulated heat stable protein 1 |
| *CBX8* | chromobox 8 |
| *CCNA2* | cyclin A2 |
| *CCNI* | cyclin I |
| *CCNJ* | cyclin J |
| *CDC20* | cell division cycle 20 |
| *CDH3* | cadherin 3 |
| *CEP170* | centrosomal protein 170 |
| *CEP57* | centrosomal protein 57 |
| *CHCHD3* | coiled-coil-helix-coiled-coil-helix domain containing 3 |
| *CHML* | CHM like, Rab escort protein 2 |
| *CHST11* | carbohydrate sulfotransferase 11 |
| *CHSY1* | chondroitin sulfate synthase 1 |
| *CIB2* | calcium and integrin binding family member 2 |
| *CKAP2* | cytoskeleton associated protein 2 |
| *CKLF* | chemokine like factor |
| *CKS2* | CDC28 protein kinase regulatory subunit 2 |
| *CNN2* | calponin 2 |
| *COL25A1* | collagen type XXV alpha 1 chain |
| *COL3A1* | collagen type III alpha 1 chain |
| *CORO1C* | coronin 1C |
| *CPNE3* | copine 3 |
| *CRK* | CRK proto-oncogene, adaptor protein |
| *CSRNP3* | cysteine and serine rich nuclear protein 3 |
| *CTCF* | CCCTC-binding factor |
| *DAAM1* | dishevelled associated activator of morphogenesis 1 |
| *DACT1* | dishevelled binding antagonist of beta catenin 1 |
| *DCAF12* | DDB1 and CUL4 associated factor 12 |
| *DCK* | deoxycytidine kinase |
| *DEPDC1B* | DEP domain containing 1B |
| *DGKD* | diacylglycerol kinase delta |
| *DLG5* | discs large MAGUK scaffold protein 5 |
| *DLGAP5* | DLG associated protein 5 |
| *DLX1* | distal-less homeobox 1 |
| *DNMT3A* | DNA methyltransferase 3 alpha |
| *DNMT3B* | DNA methyltransferase 3 beta |
| *DPEP1* | dipeptidase 1 |
| *DPYSL5* | dihydropyrimidinase like 5 |
| *DUSP12* | dual specificity phosphatase 12 |
| *DYRK2* | dual specificity tyrosine phosphorylation regulated kinase 2 |
| *E2F5* | E2F transcription factor 5 |
| *EEF1D* | eukaryotic translation elongation factor 1 delta |
| *EHD1* | EH domain containing 1 |
| *EIF4EBP2* | eukaryotic translation initiation factor 4E binding protein 2 |
| *EIF4G2* | eukaryotic translation initiation factor 4 gamma 2 |
| *ELN* | elastin |
| *EMID1* | EMI domain containing 1 |
| *EML1* | echinoderm microtubule associated protein like 1 |
| *ENO3* | enolase 3 |
| *EOMES* | eomesodermin |
| *EPB41* | erythrocyte membrane protein band 4.1 |
| *EPHA5* | EPH receptor A5 |
| *EPHA7* | EPH receptor A7 |
| *EPHB4* | EPH receptor B4 |
| *EPS8* | epidermal growth factor receptor pathway substrate 8 |
| *EZR* | ezrin |
| *FAM172A* | family with sequence similarity 172 member A |
| *FANCC* | Fanconi anemia complementation group C |
| *FCHSD2* | FCH and double SH3 domains 2 |
| *FDFT1* | farnesyl-diphosphate farnesyltransferase 1 |
| *FGD4* | FYVE, RhoGEF and PH domain containing 4 |
| *FHL3* | four and a half LIM domains 3 |
| *FOXN3* | forkhead box N3 |
| *FSCN1* | fascin actin-bundling protein 1 |
| *GINS2* | GINS complex subunit 2 |
| *GMEB1* | glucocorticoid modulatory element binding protein 1 |
| *GPM6A* | glycoprotein M6A |
| *GPR85* | G protein-coupled receptor 85 |
| *GRIN2D* | glutamate ionotropic receptor NMDA type subunit 2D |
| *GTSE1* | G2 and S-phase expressed 1 |
| *H19* | H19, imprinted maternally expressed transcript (non-protein coding) |
| *H2AFY2* | H2A histone family member Y2 |
| *HAPLN3* | hyaluronan and proteoglycan link protein 3 |
| *HCN3* | hyperpolarization activated cyclic nucleotide gated potassium channel 3 |
| *HCRTR2* | hypocretin receptor 2 |
| *HDAC2* | histone deacetylase 2 |
| *HIF1A* | hypoxia inducible factor 1 subunit alpha |
| *HIST3H2A* | histone cluster 3 H2A |
| *HMGCR* | 3-hydroxy-3-methylglutaryl-CoA reductase |
| *HMGN1* | high mobility group nucleosome binding domain 1 |
| *HSF2* | heat shock transcription factor 2 |
| *ICK* | intestinal cell kinase |
| *IDH1* | isocitrate dehydrogenase (NADP(+)) 1, cytosolic |
| *IGF2BP1* | insulin like growth factor 2 mRNA binding protein 1 |
| *IGF2BP3* | insulin like growth factor 2 mRNA binding protein 3 |
| *IGFBP5* | insulin like growth factor binding protein 5 |
| *IGFBPL1* | insulin like growth factor binding protein like 1 |
| *ING1* | inhibitor of growth family member 1 |
| *ITPA* | inosine triphosphatase |
| *ITPRIP* | inositol 1,4,5-trisphosphate receptor interacting protein |
| *KCTD12* | potassium channel tetramerization domain containing 12 |
| *KCTD15* | potassium channel tetramerization domain containing 15 |
| *KIAA1549* | KIAA1549 |
| *KIF15* | kinesin family member 15 |
| *KIF21B* | kinesin family member 21B |
| *KIF2C* | kinesin family member 2C |
| *KLF12* | Kruppel like factor 12 |
| *KLHL23* | kelch like family member 23 |
| *KLHL9* | kelch like family member 9 |
| *LAMC1* | laminin subunit gamma 1 |
| *LASP1* | LIM and SH3 protein 1 |
| *LCORL* | ligand dependent nuclear receptor corepressor like |
| *LHX2* | LIM homeobox 2 |
| *LMNB1* | lamin B1 |
| *LOXL3* | lysyl oxidase like 3 |
| *LPAR2* | lysophosphatidic acid receptor 2 |
| *LRRC40* | leucine rich repeat containing 40 |
| *LSM5* | LSM5 homolog, U6 small nuclear RNA and mRNA degradation associated |
| *MAPRE1* | microtubule associated protein RP/EB family member 1 |
| *MASP1* | mannan binding lectin serine peptidase 1 |
| *MAST2* | microtubule associated serine/threonine kinase 2 |
| *MDFI* | MyoD family inhibitor |
| *MEF2C* | myocyte enhancer factor 2C |
| *MFAP2* | microfibril associated protein 2 |
| *MGAT2* | mannosyl (alpha-1,6-)-glycoprotein beta-1,2-N-acetylglucosaminyltransferase |
| *MMP2* | matrix metallopeptidase 2 |
| *MN1* | MN1 proto-oncogene, transcriptional regulator |
| *MPG* | N-methylpurine DNA glycosylase |
| *MSI1* | musashi RNA binding protein 1 |
| *MTA2* | metastasis associated 1 family member 2 |
| *MTSS1* | MTSS1, I-BAR domain containing |
| *MYNN* | myoneurin |
| *NBEAL1* | neurobeachin like 1 |
| *NCOA6* | nuclear receptor coactivator 6 |
| *NDUFA11* | NADH:ubiquinone oxidoreductase subunit A11 |
| *NEDD4L* | neural precursor cell expressed, developmentally down-regulated 4-like, E3 ubiquitin protein ligase |
| *NGFR* | nerve growth factor receptor |
| *NID1* | nidogen 1 |
| *NMU* | neuromedin U |
| *NPM1* | nucleophosmin 1 |
| *NRAS* | NRAS proto-oncogene, GTPase |
| *NUP62* | nucleoporin 62 |
| *NXPH4* | neurexophilin 4 |
| *ODC1* | ornithine decarboxylase 1 |
| *PACS1* | phosphofurin acidic cluster sorting protein 1 |
| *PAK2* | p21 (RAC1) activated kinase 2 |
| *PANK1* | pantothenate kinase 1 |
| *PBRM1* | polybromo 1 |
| *PDCL* | phosducin like |
| *PDE10A* | phosphodiesterase 10A |
| *PDE4D* | phosphodiesterase 4D |
| *PDGFC* | platelet derived growth factor C |
| *PHF2* | PHD finger protein 2 |
| *PIK3R3* | phosphoinositide-3-kinase regulatory subunit 3 |
| *PITPNC1* | phosphatidylinositol transfer protein cytoplasmic 1 |
| *PLSCR3* | phospholipid scramblase 3 |
| *PMF1* | polyamine modulated factor 1 |
| *POLD1* | DNA polymerase delta 1, catalytic subunit |
| *POLDIP3* | DNA polymerase delta interacting protein 3 |
| *POLR2F* | RNA polymerase II subunit F |
| *PPIH* | peptidylprolyl isomerase H |
| *PPP1R14B* | protein phosphatase 1 regulatory inhibitor subunit 14B |
| *PPP1R35* | protein phosphatase 1 regulatory subunit 35 |
| *PPP2R3A* | protein phosphatase 2 regulatory subunit B''alpha |
| *PPP2R5E* | protein phosphatase 2 regulatory subunit B'epsilon |
| *PPP3CC* | protein phosphatase 3 catalytic subunit gamma |
| *PPP4C* | protein phosphatase 4 catalytic subunit |
| *PRDM8* | PR/SET domain 8 |
| *PRKAR2B* | protein kinase cAMP-dependent type II regulatory subunit beta |
| *PRPF40A* | pre-mRNA processing factor 40 homolog A |
| *PRR5* | proline rich 5 |
| *PTGFRN* | prostaglandin F2 receptor inhibitor |
| *PTPRS* | protein tyrosine phosphatase, receptor type S |
| *PTTG1IP* | PTTG1 interacting protein |
| *PTX3* | pentraxin 3 |
| *QARS* | glutaminyl-tRNA synthetase |
| *RAB10* | RAB10, member RAS oncogene family |
| *RALY* | RALY heterogeneous nuclear ribonucleoprotein |
| *RASSF1* | Ras association domain family member 1 |
| *RAVER1* | ribonucleoprotein, PTB binding 1 |
| *RBFOX2* | RNA binding fox-1 homolog 2 |
| *RBP1* | retinol binding protein 1 |
| *RCC1* | regulator of chromosome condensation 1 |
| *RELN* | reelin |
| *RFC4* | replication factor C subunit 4 |
| *RFX3* | regulatory factor X3 |
| *RLF* | rearranged L-myc fusion |
| *RNF122* | ring finger protein 122 |
| *RNF182* | ring finger protein 182 |
| *RNF2* | ring finger protein 2 |
| *ROBO1* | roundabout guidance receptor 1 |
| *RPL11* | ribosomal protein L11 |
| *RPL35* | ribosomal protein L35 |
| *RPS10* | ribosomal protein S10 |
| *RPS24* | ribosomal protein S24 |
| *RPS25* | ribosomal protein S25 |
| *RPS3* | ribosomal protein S3 |
| *RPS6* | ribosomal protein S6 |
| *RRP1B* | ribosomal RNA processing 1B |
| *RSBN1* | round spermatid basic protein 1 |
| *RUVBL1* | RuvB like AAA ATPase 1 |
| *SARNP* | SAP domain containing ribonucleoprotein |
| *SCN3A* | sodium voltage-gated channel alpha subunit 3 |
| *SCRT2* | scratch family transcriptional repressor 2 |
| *SCYL2* | SCY1 like pseudokinase 2 |
| *SDC1* | syndecan 1 |
| *SDK1* | sidekick cell adhesion molecule 1 |
| *SEC11A* | SEC11 homolog A, signal peptidase complex subunit |
| *SEC61A1* | Sec61 translocon alpha 1 subunit |
| *SEMA6C* | semaphorin 6C |
| *SENP1* | SUMO specific peptidase 1 |
| *SENP7* | SUMO specific peptidase 7 |
| *SEZ6* | seizure related 6 homolog |
| *SFRP1* | secreted frizzled related protein 1 |
| *SH3PXD2B* | SH3 and PX domains 2B |
| *SKA2* | spindle and kinetochore associated complex subunit 2 |
| *SLC16A9* | solute carrier family 16 member 9 |
| *SLC1A5* | solute carrier family 1 member 5 |
| *SLC25A24* | solute carrier family 25 member 24 |
| *SLC35F2* | solute carrier family 35 member F2 |
| *SMAD4* | SMAD family member 4 |
| *SMARCD1* | SWI/SNF related, matrix associated, actin dependent regulator of chromatin, subfamily d, member 1 |
| *SMO* | smoothened, frizzled class receptor |
| *SMPD3* | sphingomyelin phosphodiesterase 3 |
| *SP3* | Sp3 transcription factor |
| *SRD5A1* | steroid 5 alpha-reductase 1 |
| *SS18L1* | SS18L1, nBAF chromatin remodeling complex subunit |
| *ST3GAL2* | ST3 beta-galactoside alpha-2,3-sialyltransferase 2 |
| *ST6GAL2* | ST6 beta-galactoside alpha-2,6-sialyltransferase 2 |
| *STC1* | stanniocalcin 1 |
| *STK32B* | serine/threonine kinase 32B |
| *STK38* | serine/threonine kinase 38 |
| *STMN1* | stathmin 1 |
| *STYX* | serine/threonine/tyrosine interacting protein |
| *SUFU* | SUFU negative regulator of hedgehog signaling |
| *SUMO2* | small ubiquitin-like modifier 2 |
| *SUV39H2* | suppressor of variegation 3-9 homolog 2 |
| *SYNC* | syncoilin, intermediate filament protein |
| *SYT6* | synaptotagmin 6 |
| *TACC3* | transforming acidic coiled-coil containing protein 3 |
| *TBC1D10A* | TBC1 domain family member 10A |
| *TCF3* | transcription factor 3 |
| *TFAP2C* | transcription factor AP-2 gamma |
| *TLE3* | transducin like enhancer of split 3 |
| *TMED9* | transmembrane p24 trafficking protein 9 |
| *TMEFF1* | transmembrane protein with EGF like and two follistatin like domains 1 |
| *TMPO* | thymopoietin |
| *TNPO1* | transportin 1 |
| *TNRC18* | trinucleotide repeat containing 18 |
| *TPRKB* | TP53RK binding protein |
| *TPX2* | TPX2, microtubule nucleation factor |
| *TRIM16* | tripartite motif containing 16 |
| *TRIM24* | tripartite motif containing 24 |
| *TRIM36* | tripartite motif containing 36 |
| *TRIOBP* | TRIO and F-actin binding protein |
| *TTK* | TTK protein kinase |
| *UBA52* | ubiquitin A-52 residue ribosomal protein fusion product 1 |
| *UCP2* | uncoupling protein 2 |
| *UHRF1* | ubiquitin like with PHD and ring finger domains 1 |
| *USF1* | upstream transcription factor 1 |
| *USP42* | ubiquitin specific peptidase 42 |
| *VCAN* | versican |
| *WNT5B* | Wnt family member 5B |
| *WNT7A* | Wnt family member 7A |
| *XPR1* | xenotropic and polytropic retrovirus receptor 1 |
| *YPEL1* | yippee like 1 |
| *YTHDF2* | YTH N6-methyladenosine RNA binding protein 2 |
| *YWHAE* | tyrosine 3-monooxygenase/tryptophan 5-monooxygenase activation protein epsilon |
| *ZBTB10* | zinc finger and BTB domain containing 10 |
| *ZBTB5* | zinc finger and BTB domain containing 5 |
| *ZCCHC14* | zinc finger CCHC-type containing 14 |
| *ZIK1* | zinc finger protein interacting with K protein 1 |
| *ZNF85* | zinc finger protein 85 |
| *ZSWIM4* | zinc finger SWIM-type containing 4 |
| *ZSWIM5* | zinc finger SWIM-type containing 5 |

Table S2.

Study participants’ characteristics correlation with mesocorticolimbic ePRS-IR

| **Sample descriptives** | **Spearman correlation coefficient** | **P-value** |
| --- | --- | --- |
| **MAVAN cohort** | | |
| Gender ^a^ | NA | 0.785 |
| Birth weight (grams)^c^ | -0.05 | 0.459 |
| Gestational age (weeks)^c^ | 0.02 | 0.817 |
| Family income below Low Income Cut Off(Canada 2005) ^a^ | NA | 0.945 |
| **SAGE cohort** | | |
| Gender ^a^ | NA | 0.193 |
| Family income below $20K ^a^ | NA | 0.266 |
| Age at interview^c^ | 0.03 | 0.159 |
| Study source ^b^ | NA | 0.033* |

Study participants’ characteristics correlation with hippocampal ePRS-IR

| **Sample descriptives** | **Spearman correlation coefficient** | **P-value** |
| --- | --- | --- |
| **MAVAN cohort** | | |
| Gender ^a^ | NA | 0.584 |
| Birth weight (grams)^c^ | -0.07 | 0.276 |
| Gestational age (weeks)^c^ | -0.04 | 0.585 |
| Family income below Low Income Cut Off(Canada 2005) ^a^ | NA | 0.063 |
| **ADA cohort** | | |
| Gender ^a^ | NA | 0.558 |
| Study site ^b^ | NA | 0.180 |
| Age at onset^c^ | 0.08 | 0.033* |

Statistical analyses: ^a^ t-tests, ^b^ one-way ANOVA, ^c^ Spearman correlation

Table S3.

List of genes selected for composing the hippocampal ePRS-IR (see details in the text).

| ***Gene Symbol*** | **Description** |
| --- | --- |
| *ADGRG1* | adhesion G protein-coupled receptor G1 |
| *AGO1* | argonaute 1, RISC catalytic component |
| *ALYREF* | Aly/REF export factor |
| *B4GALT5* | beta-1,4-galactosyltransferase 5 |
| *BBC3* | BCL2 binding component 3 |
| *BCCIP* | BRCA2 and CDKN1A interacting protein |
| *BCHE* | butyrylcholinesterase |
| *BCL11B* | B cell CLL/lymphoma 11B |
| *BCL7C* | BCL tumor suppressor 7C |
| *BCL9* | B cell CLL/lymphoma 9 |
| *BCLAF1* | BCL2 associated transcription factor 1 |
| *BFAR* | bifunctional apoptosis regulator |
| *BIRC2* | baculoviral IAP repeat containing 2 |
| *BOC* | BOC cell adhesion associated, oncogene regulated |
| *BPTF* | bromodomain PHD finger transcription factor |
| *BRD1* | bromodomain containing 1 |
| *BRIP1* | BRCA1 interacting protein C-terminal helicase 1 |
| *BTF3* | basic transcription factor 3 |
| *BTG1* | BTG anti-proliferation factor 1 |
| *BZW1* | basic leucine zipper and W2 domains 1 |
| *BZW2* | basic leucine zipper and W2 domains 2 |
| *CADM1* | cell adhesion molecule 1 |
| *CBLB* | Cbl proto-oncogene B |
| *CBX2* | chromobox 2 |
| *CBX3* | chromobox 3 |
| *CBX5* | chromobox 5 |
| *CCDC102A* | coiled-coil domain containing 102A |
| *CCDC138* | coiled-coil domain containing 138 |
| *CCDC28B* | coiled-coil domain containing 28B |
| *CCDC40* | coiled-coil domain containing 40 |
| *CCDC97* | coiled-coil domain containing 97 |
| *CCNF* | cyclin F |
| *CCSER1* | coiled-coil serine rich protein 1 |
| *CCT2* | chaperonin containing TCP1 subunit 2 |
| *CCT3* | chaperonin containing TCP1 subunit 3 |
| *CD248* | CD248 molecule |
| *CD93* | CD93 molecule |
| *CDC25A* | cell division cycle 25A |
| *CDC42* | cell division cycle 42 |
| *CDCA5* | cell division cycle associated 5 |
| *CDCA8* | cell division cycle associated 8 |
| *CDH11* | cadherin 11 |
| *CDK1* | cyclin dependent kinase 1 |
| *CDKN1B* | cyclin dependent kinase inhibitor 1B |
| *CDKN2C* | cyclin dependent kinase inhibitor 2C |
| *CDON* | cell adhesion associated, oncogene regulated |
| *CELSR3* | cadherin EGF LAG seven-pass G-type receptor 3 |
| *CENPF* | centromere protein F |
| *CENPK* | centromere protein K |
| *CHODL* | chondrolectin |
| *CHST10* | carbohydrate sulfotransferase 10 |
| *CHST14* | carbohydrate sulfotransferase 14 |
| *CKLF* | chemokine like factor |
| *CKS2* | CDC28 protein kinase regulatory subunit 2 |
| *CNOT1* | CCR4-NOT transcription complex subunit 1 |
| *CNOT2* | CCR4-NOT transcription complex subunit 2 |
| *CNOT6* | CCR4-NOT transcription complex subunit 6 |
| *CNOT6L* | CCR4-NOT transcription complex subunit 6 like |
| *CNOT8* | CCR4-NOT transcription complex subunit 8 |
| *CNR1* | cannabinoid receptor 1 |
| *COIL* | coilin |
| *COL11A1* | collagen type XI alpha 1 chain |
| *COL2A1* | collagen type II alpha 1 chain |
| *COL3A1* | collagen type III alpha 1 chain |
| *COLGALT1* | collagen beta(1-O)galactosyltransferase 1 |
| *CPXM1* | carboxypeptidase X, M14 family member 1 |
| *CRB1* | crumbs 1, cell polarity complex component |
| *CREB3L4* | cAMP responsive element binding protein 3 like 4 |
| *CREBBP* | CREB binding protein |
| *CRY1* | cryptochrome circadian regulator 1 |
| *CSNK1G1* | casein kinase 1 gamma 1 |
| *CTBP2* | C-terminal binding protein 2 |
| *CTDSPL2* | CTD small phosphatase like 2 |
| *CTNNB1* | catenin beta 1 |
| *CUL7* | cullin 7 |
| *CUX1* | cut like homeobox 1 |
| *CYBA* | cytochrome b-245 alpha chain |
| *CYFIP1* | cytoplasmic FMR1 interacting protein 1 |
| *CYR61* | cysteine rich angiogenic inducer 61 |
| *DAAM1* | dishevelled associated activator of morphogenesis 1 |
| *DACH1* | dachshund family transcription factor 1 |
| *DAG1* | dystroglycan 1 |
| *DCHS1* | dachsous cadherin-related 1 |
| *DDIT4* | DNA damage inducible transcript 4 |
| *DDX21* | DExD-box helicase 21 |
| *DEK* | DEK proto-oncogene |
| *DEPDC1B* | DEP domain containing 1B |
| *DGKD* | diacylglycerol kinase delta |
| *DHTKD1* | dehydrogenase E1 and transketolase domain containing 1 |
| *DHX40* | DEAH-box helicase 40 |
| *DLGAP5* | DLG associated protein 5 |
| *DNAJB5* | DnaJ heat shock protein family (Hsp40) member B5 |
| *DNALI1* | dynein axonemal light intermediate chain 1 |
| *DNMBP* | dynamin binding protein |
| *DPH5* | diphthamide biosynthesis 5 |
| *DSG2* | desmoglein 2 |
| *DTD1* | D-tyrosyl-tRNA deacylase 1 |
| *DUSP18* | dual specificity phosphatase 18 |
| *DUSP4* | dual specificity phosphatase 4 |
| *DVL3* | dishevelled segment polarity protein 3 |
| *DYM* | dymeclin |
| *E2F2* | E2F transcription factor 2 |
| *E2F7* | E2F transcription factor 7 |
| *EBPL* | emopamil binding protein like |
| *ECT2* | epithelial cell transforming 2 |
| *EDNRA* | endothelin receptor type A |
| *EEF1G* | eukaryotic translation elongation factor 1 gamma |
| *EEFSEC* | eukaryotic elongation factor, selenocysteine-tRNA specific |
| *EHMT1* | euchromatic histone lysine methyltransferase 1 |
| *EHMT2* | euchromatic histone lysine methyltransferase 2 |
| *EIF3C* | eukaryotic translation initiation factor 3 subunit C |
| *EIF3D* | eukaryotic translation initiation factor 3 subunit D |
| *EIF3H* | eukaryotic translation initiation factor 3 subunit H |
| *EIF4B* | eukaryotic translation initiation factor 4B |
| *EIF4G2* | eukaryotic translation initiation factor 4 gamma 2 |
| *ELF2* | E74 like ETS transcription factor 2 |
| *ELOVL2* | ELOVL fatty acid elongase 2 |
| *ENTPD7* | ectonucleoside triphosphate diphosphohydrolase 7 |
| *EPB41* | erythrocyte membrane protein band 4.1 |
| *EPHA7* | EPH receptor A7 |
| *EVC* | EvC ciliary complex subunit 1 |
| *EXOSC8* | exosome component 8 |
| *EXT2* | exostosin glycosyltransferase 2 |
| *EZH2* | enhancer of zeste 2 polycomb repressive complex 2 subunit |
| *FABP7* | fatty acid binding protein 7 |
| *FBLIM1* | filamin binding LIM protein 1 |
| *FBXO46* | F-box protein 46 |
| *FBXW8* | F-box and WD repeat domain containing 8 |
| *FDFT1* | farnesyl-diphosphate farnesyltransferase 1 |
| *FEM1C* | fem-1 homolog C |
| *FGD3* | FYVE, RhoGEF and PH domain containing 3 |
| *FGD4* | FYVE, RhoGEF and PH domain containing 4 |
| *FKBP10* | FK506 binding protein 10 |
| *FOXG1* | forkhead box G1 |
| *FOXM1* | forkhead box M1 |
| *FOXN3* | forkhead box N3 |
| *FOXN4* | forkhead box N4 |
| *FREM2* | FRAS1 related extracellular matrix protein 2 |
| *FRMD3* | FERM domain containing 3 |
| *FRMD4A* | FERM domain containing 4A |
| *FTSJ3* | FtsJ RNA methyltransferase homolog 3 |
| *FYN* | FYN proto-oncogene, Src family tyrosine kinase |
| *G3BP1* | G3BP stress granule assembly factor 1 |
| *GADD45A* | growth arrest and DNA damage inducible alpha |
| *GAL3ST4* | galactose-3-O-sulfotransferase 4 |
| *GART* | phosphoribosylglycinamide formyltransferase, |
| *GCNT2* | glucosaminyl (N-acetyl) transferase 2, I-branching enzyme |
| *GDI2* | GDP dissociation inhibitor 2 |
| *GEMIN5* | gem nuclear organelle associated protein 5 |
| *GFPT1* | glutamine--fructose-6-phosphate transaminase 1 |
| *GLI2* | GLI family zinc finger 2 |
| *GLI3* | GLI family zinc finger 3 |
| *GMEB1* | glucocorticoid modulatory element binding protein 1 |
| *GNAI3* | G protein subunit alpha i3 |
| *GNL2* | G protein nucleolar 2 |
| *GPC1* | glypican 1 |
| *GPR139* | G protein-coupled receptor 139 |
| *GRIP1* | glutamate receptor interacting protein 1 |
| *GSTCD* | glutathione S-transferase C-terminal domain containing |
| *GTF2IRD1* | GTF2I repeat domain containing 1 |
| *GTSE1* | G2 and S-phase expressed 1 |
| *H19* | H19, imprinted maternally expressed transcript (non-protein coding) |
| *H2AFY2* | H2A histone family member Y2 |
| *H3F3A* | H3 histone family member 3A |
| *H3F3B* | H3 histone family member 3B |
| *HAPLN3* | hyaluronan and proteoglycan link protein 3 |
| *HDAC2* | histone deacetylase 2 |
| *HDGF* | heparin binding growth factor |
| *HDHD5* | haloacid dehalogenase like hydrolase domain containing 5 |
| *HEATR6* | HEAT repeat containing 6 |
| *HEBP2* | heme binding protein 2 |
| *HECW1* | HECT, C2 and WW domain containing E3 ubiquitin protein ligase 1 |
| *HIF1A* | hypoxia inducible factor 1 alpha subunit |
| *HMGA2* | high mobility group AT-hook 2 |
| *HMGB1* | high mobility group box 1 |
| *HNRNPA1* | heterogeneous nuclear ribonucleoprotein A1 |
| *HNRNPR* | heterogeneous nuclear ribonucleoprotein R |
| *HNRNPU* | heterogeneous nuclear ribonucleoprotein U |
| *HSF2* | heat shock transcription factor 2 |
| *ID4* | inhibitor of DNA binding 4, HLH protein |
| *IFT140* | intraflagellar transport 140 |
| *IGF2BP3* | insulin like growth factor 2 mRNA binding protein 3 |
| *IGSF9* | immunoglobulin superfamily member 9 |
| *ING1* | inhibitor of growth family member 1 |
| *ING4* | inhibitor of growth family member 4 |
| *IPO4* | importin 4 |
| *IQGAP2* | IQ motif containing GTPase activating protein 2 |
| *IQGAP3* | IQ motif containing GTPase activating protein 3 |
| *IRF2BP2* | interferon regulatory factor 2 binding protein 2 |
| *IRS1* | insulin receptor substrate 1 |
| *ITGA6* | integrin subunit alpha 6 |
| *ITGB8* | integrin subunit beta 8 |
| *JAKMIP2* | janus kinase and microtubule interacting protein 2 |
| *JAM2* | junctional adhesion molecule 2 |
| *KCNK10* | potassium two pore domain channel subfamily K member 10 |
| *KDM4A* | lysine demethylase 4A |
| *KIDINS220* | kinase D interacting substrate 220 |
| *KIF11* | kinesin family member 11 |
| *KIF14* | kinesin family member 14 |
| *KIF15* | kinesin family member 15 |
| *KIF18A* | kinesin family member 18A |
| *KIF22* | kinesin family member 22 |
| *KIF23* | kinesin family member 23 |
| *KIFC1* | kinesin family member C1 |
| *KLF12* | Kruppel like factor 12 |
| *KLF6* | Kruppel like factor 6 |
| *KLHDC8A* | kelch domain containing 8A |
| *KLHL7* | kelch like family member 7 |
| *KNL1* | kinetochore scaffold 1 |
| *KPNA2* | karyopherin subunit alpha 2 |
| *KPNB1* | karyopherin subunit beta 1 |
| *LIMD1* | LIM domains containing 1 |
| *LIN7C* | lin-7 homolog C, crumbs cell polarity complex component |
| *LIN9* | lin-9 DREAM MuvB core complex component |
| *LMAN2L* | lectin, mannose binding 2 like |
| *LMNB1* | lamin B1 |
| *LOXL1* | lysyl oxidase like 1 |
| *LOXL2* | lysyl oxidase like 2 |
| *LOXL3* | lysyl oxidase like 3 |
| *LPAR2* | lysophosphatidic acid receptor 2 |
| *LRP5* | LDL receptor related protein 5 |
| *LRP8* | LDL receptor related protein 8 |
| *LRRC1* | leucine rich repeat containing 1 |
| *LRRC17* | leucine rich repeat containing 17 |
| *LRRC61* | leucine rich repeat containing 61 |
| *LRRN1* | leucine rich repeat neuronal 1 |
| *LTBP1* | latent transforming growth factor beta binding protein 1 |
| *LYAR* | Ly1 antibody reactive |
| *LYN* | LYN proto-oncogene, Src family tyrosine kinase |
| *MAD2L2* | mitotic arrest deficient 2 like 2 |
| *MAN2B1* | mannosidase alpha class 2B member 1 |
| *MAPK8* | mitogen-activated protein kinase 8 |
| *MASP1* | mannan binding lectin serine peptidase 1 |
| *MBTD1* | mbt domain containing 1 |
| *MCM4* | minichromosome maintenance complex component 4 |
| *MCUB* | mitochondrial calcium uniporter dominant negative B subunit |
| *MDM1* | Mdm1 nuclear protein |
| *MED1* | mediator complex subunit 1 |
| *MED20* | mediator complex subunit 20 |
| *MELK* | maternal embryonic leucine zipper kinase |
| *METAP2* | methionyl aminopeptidase 2 |
| *MFNG* | MFNG O-fucosylpeptide 3-beta-N-acetylglucosaminyltransferase |
| *MLLT3* | MLLT3, super elongation complex subunit |
| *MMP14* | matrix metallopeptidase 14 |
| *MMP16* | matrix metallopeptidase 16 |
| *MMP25* | matrix metallopeptidase 25 |
| *MRC2* | mannose receptor C type 2 |
| *MSH2* | mutS homolog 2 |
| *MSI1* | musashi RNA binding protein 1 |
| *MSL1* | male specific lethal 1 homolog |
| *MTSS1* | MTSS1, I-BAR domain containing |
| *MYCL* | MYCL proto-oncogene, bHLH transcription factor |
| *MYO3A* | myosin IIIA |
| *NAB1* | NGFI-A binding protein 1 |
| *NACA* | nascent polypeptide-associated complex alpha subunit |
| *NBEAL1* | neurobeachin like 1 |
| *NCAPD2* | non-SMC condensin I complex subunit D2 |
| *NCAPG* | non-SMC condensin I complex subunit G |
| *NCOA6* | nuclear receptor coactivator 6 |
| *NECTIN3* | nectin cell adhesion molecule 3 |
| *NEIL3* | nei like DNA glycosylase 3 |
| *NEK2* | NIMA related kinase 2 |
| *NELL1* | neural EGFL like 1 |
| *NES* | nestin |
| *NFIA* | nuclear factor I A |
| *NFIB* | nuclear factor I B |
| *NHSL1* | NHS like 1 |
| *NLGN1* | neuroligin 1 |
| *NMU* | neuromedin U |
| *NOP53* | NOP53 ribosome biogenesis factor |
| *NPR3* | natriuretic peptide receptor 3 |
| *NR2E1* | nuclear receptor subfamily 2 group E member 1 |
| *NRF1* | nuclear respiratory factor 1 |
| *NSD1* | nuclear receptor binding SET domain protein 1 |
| *NSG1* | neuronal vesicle trafficking associated 1 |
| *NUF2* | NUF2, NDC80 kinetochore complex component |
| *NUP107* | nucleoporin 107 |
| *NUP205* | nucleoporin 205 |
| *NUP62* | nucleoporin 62 |
| *NUSAP1* | nucleolar and spindle associated protein 1 |
| *NYNRIN* | NYN domain and retroviral integrase containing |
| *OSBPL11* | oxysterol binding protein like 11 |
| *OSBPL5* | oxysterol binding protein like 5 |
| *PA2G4* | proliferation-associated 2G4 |
| *PABPC1* | poly(A) binding protein cytoplasmic 1 |
| *PABPC3* | poly(A) binding protein cytoplasmic 3 |
| *PAICS* | phosphoribosylaminoimidazole carboxylase |
| *PANK1* | pantothenate kinase 1 |
| *PANK3* | pantothenate kinase 3 |
| *PANX1* | pannexin 1 |
| *PARD3B* | par-3 family cell polarity regulator beta |
| *PATZ1* | POZ/BTB and AT hook containing zinc finger 1 |
| *PAWR* | pro-apoptotic WT1 regulator |
| *PAX6* | paired box 6 |
| *PBRM1* | polybromo 1 |
| *PBX1* | PBX homeobox 1 |
| *PCBP2* | poly(rC) binding protein 2 |
| *PCDHB18P* | protocadherin beta 18 pseudogene |
| *PCIF1* | PDX1 C-terminal inhibiting factor 1 |
| *PCNA* | proliferating cell nuclear antigen |
| *PDE10A* | phosphodiesterase 10A |
| *PDE7A* | phosphodiesterase 7A |
| *PDGFC* | platelet derived growth factor C |
| *PDIA6* | protein disulfide isomerase family A member 6 |
| *PDIK1L* | PDLIM1 interacting kinase 1 like |
| *PELI1* | pellino E3 ubiquitin protein ligase 1 |
| *PELI2* | pellino E3 ubiquitin protein ligase family member 2 |
| *PFAS* | phosphoribosylformylglycinamidine synthase |
| *PGM2* | phosphoglucomutase 2 |
| *PHF20* | PHD finger protein 20 |
| *PHF21A* | PHD finger protein 21A |
| *PHIP* | pleckstrin homology domain interacting protein |
| *PHTF2* | putative homeodomain transcription factor 2 |
| *PLEKHG2* | pleckstrin homology and RhoGEF domain containing G2 |
| *PLK4* | polo like kinase 4 |
| *PNN* | pinin, desmosome associated protein |
| *POLE* | DNA polymerase epsilon, catalytic subunit |
| *POLE2* | DNA polymerase epsilon 2, accessory subunit |
| *POLR1E* | RNA polymerase I subunit E |
| *POLR3D* | RNA polymerase III subunit D |
| *PPM1D* | protein phosphatase, Mg2+/Mn2+ dependent 1D |
| *PPP1CC* | protein phosphatase 1 catalytic subunit gamma |
| *PPP1R14C* | protein phosphatase 1 regulatory inhibitor subunit 14C |
| *PPP1R18* | protein phosphatase 1 regulatory subunit 18 |
| *PPP2R5E* | protein phosphatase 2 regulatory subunit B'epsilon |
| *PPP4R1* | protein phosphatase 4 regulatory subunit 1 |
| *PPP4R3A* | protein phosphatase 4 regulatory subunit 3A |
| *PPP4R3B* | protein phosphatase 4 regulatory subunit 3B |
| *PRC1* | protein regulator of cytokinesis 1 |
| *PRCC* | papillary renal cell carcinoma (translocation-associated) |
| *PRKAR2A* | protein kinase cAMP-dependent type II regulatory subunit alpha |
| *PRMT5* | protein arginine methyltransferase 5 |
| *PROM1* | prominin 1 |
| *PRPF38A* | pre-mRNA processing factor 38A |
| *PRPF40A* | pre-mRNA processing factor 40 homolog A |
| *PRPSAP2* | phosphoribosyl pyrophosphate synthetase associated protein 2 |
| *PRR14* | proline rich 14 |
| *PTGFRN* | prostaglandin F2 receptor inhibitor |
| *PTK7* | protein tyrosine kinase 7 (inactive) |
| *PTMA* | prothymosin alpha |
| *PTN* | pleiotrophin |
| *PTPN12* | protein tyrosine phosphatase, non-receptor type 12 |
| *PTPN9* | protein tyrosine phosphatase, non-receptor type 9 |
| *PTPRZ1* | protein tyrosine phosphatase, receptor type Z1 |
| *PTTG1IP* | PTTG1 interacting protein |
| *PXDN* | peroxidasin |
| *RAB10* | RAB10, member RAS oncogene family |
| *RAB13* | RAB13, member RAS oncogene family |
| *RAB8A* | RAB8A, member RAS oncogene family |
| *RACGAP1* | Rac GTPase activating protein 1 |
| *RACK1* | receptor for activated C kinase 1 |
| *RAD18* | RAD18, E3 ubiquitin protein ligase |
| *RAD51AP1* | RAD51 associated protein 1 |
| *RARA* | retinoic acid receptor alpha |
| *RARS2* | arginyl-tRNA synthetase 2, mitochondrial |
| *RBFOX2* | RNA binding fox-1 homolog 2 |
| *RBM4B* | RNA binding motif protein 4B |
| *RCN1* | reticulocalbin 1 |
| *REC8* | REC8 meiotic recombination protein |
| *RECQL* | RecQ like helicase |
| *RELN* | reelin |
| *REST* | RE1 silencing transcription factor |
| *REV3L* | REV3 like, DNA directed polymerase zeta catalytic subunit |
| *RFX1* | regulatory factor X1 |
| *RFX3* | regulatory factor X3 |
| *RFX4* | regulatory factor X4 |
| *RFX7* | regulatory factor X7 |
| *RHOBTB1* | Rho related BTB domain containing 1 |
| *RIOK1* | RIO kinase 1 |
| *RNF165* | ring finger protein 165 |
| *ROBO1* | roundabout guidance receptor 1 |
| *RPL17* | ribosomal protein L17 |
| *RPL18A* | ribosomal protein L18a |
| *RPL19* | ribosomal protein L19 |
| *RPL22L1* | ribosomal protein L22 like 1 |
| *RPL27A* | ribosomal protein L27a |
| *RPL30* | ribosomal protein L30 |
| *RPL32* | ribosomal protein L32 |
| *RPL35A* | ribosomal protein L35a |
| *RPL5* | ribosomal protein L5 |
| *RPL6* | ribosomal protein L6 |
| *RPLP2* | ribosomal protein lateral stalk subunit P2 |
| *RPS29* | ribosomal protein S29 |
| *RPS3* | ribosomal protein S3 |
| *RPS3A* | ribosomal protein S3A |
| *RPS6* | ribosomal protein S6 |
| *RPSA* | ribosomal protein SA |
| *RRAS2* | RAS related 2 |
| *RRN3* | RRN3 homolog, RNA polymerase I transcription factor |
| *RSPRY1* | ring finger and SPRY domain containing 1 |
| *RUNX1T1* | RUNX1 translocation partner 1 |
| *SAAL1* | serum amyloid A like 1 |
| *SAFB* | scaffold attachment factor B |
| *SALL3* | spalt like transcription factor 3 |
| *SEC61B* | Sec61 translocon beta subunit |
| *SEMA4C* | semaphorin 4C |
| *SEMA4G* | semaphorin 4G |
| *SEMA5B* | semaphorin 5B |
| *SENP1* | SUMO1/sentrin specific peptidase 1 |
| *SERBP1* | SERPINE1 mRNA binding protein 1 |
| *SERINC2* | serine incorporator 2 |
| *SESTD1* | SEC14 and spectrin domain containing 1 |
| *SET* | SET nuclear proto-oncogene |
| *SF1* | splicing factor 1 |
| *SF3B3* | splicing factor 3b subunit 3 |
| *SFT2D3* | SFT2 domain containing 3 |
| *SGF29* | SAGA complex associated factor 29 |
| *SGO2* | shugoshin 2 |
| *SH3PXD2B* | SH3 and PX domains 2B |
| *SHCBP1* | SHC binding and spindle associated 1 |
| *SIN3A* | SIN3 transcription regulator family member A |
| *SIX5* | SIX homeobox 5 |
| *SKIL* | SKI like proto-oncogene |
| *SLC29A4* | solute carrier family 29 member 4 |
| *SLC2A10* | solute carrier family 2 member 10 |
| *SLC39A1* | solute carrier family 39 member 1 |
| *SLC43A2* | solute carrier family 43 member 2 |
| *SLC43A2* | solute carrier family 43 member 2 |
| *SLC43A3* | solute carrier family 43 member 3 |
| *SLC6A11* | solute carrier family 6 member 11 |
| *SMARCA5* | SWI/SNF related, matrix associated, actin dependent  regulator of chromatin, subfamily a, member 5 |
| *SMARCC1* | SWI/SNF related, matrix associated, actin dependent  regulator of chromatin subfamily c member 1 |
| *SMC5* | structural maintenance of chromosomes 5 |
| *SMCHD1* | structural maintenance of chromosomes flexible hinge  domain containing 1 |
| *SMO* | smoothened, frizzled class receptor |
| *SMTNL2* | smoothelin like 2 |
| *SNAI2* | snail family transcriptional repressor 2 |
| *SNX7* | sorting nexin 7 |
| *SOCS1* | suppressor of cytokine signaling 1 |
| *SOX11* | SRY-box 11 |
| *SOX2* | SRY-box 2 |
| *SOX4* | SRY-box 4 |
| *SOX5* | SRY-box 5 |
| *SOX6* | SRY-box 6 |
| *SPAG5* | sperm associated antigen 5 |
| *SPAST* | spastin |
| *SPECC1L* | sperm antigen with calponin homology and coiled-coil domains 1 like |
| *SPINK8* | serine peptidase inhibitor, Kazal type 8 (putative) |
| *SPRY1* | sprouty RTK signaling antagonist 1 |
| *SRBD1* | S1 RNA binding domain 1 |
| *SRC* | SRC proto-oncogene, non-receptor tyrosine kinase |
| *SRGAP2* | SLIT-ROBO Rho GTPase activating protein 2 |
| *SRR* | serine racemase |
| *ST6GAL2* | ST6 beta-galactoside alpha-2,6-sialyltransferase 2 |
| *ST7* | suppression of tumorigenicity 7 |
| *ST8SIA4* | ST8 alpha-N-acetyl-neuraminide alpha-2,8-sialyltransferase 4 |
| *STAG1* | stromal antigen 1 |
| *STK38* | serine/threonine kinase 38 |
| *STMN1* | stathmin 1 |
| *SUFU* | SUFU negative regulator of hedgehog signaling |
| *SVIL* | supervillin |
| *SYNC* | syncoilin, intermediate filament protein |
| *SYNCRIP* | synaptotagmin binding cytoplasmic RNA interacting protein |
| *TAF15* | TATA-box binding protein associated factor 15 |
| *TARS* | threonyl-tRNA synthetase |
| *TCF12* | transcription factor 12 |
| *TDG* | thymine DNA glycosylase |
| *TDRD3* | tudor domain containing 3 |
| *TEAD1* | TEA domain transcription factor 1 |
| *TGIF2* | TGFB induced factor homeobox 2 |
| *TGS1* | trimethylguanosine synthase 1 |
| *THRAP3* | thyroid hormone receptor associated protein 3 |
| *TIA1* | TIA1 cytotoxic granule associated RNA binding protein |
| *TK1* | thymidine kinase 1 |
| *TMEFF1* | transmembrane protein with EGF like and two follistatin  like domains 1 |
| *TMEM97* | transmembrane protein 97 |
| *TMPO* | thymopoietin |
| *TMTC4* | transmembrane and tetratricopeptide repeat containing 4 |
| *TNC* | tenascin C |
| *TNRC18* | trinucleotide repeat containing 18 |
| *TOP2A* | DNA topoisomerase II alpha |
| *TOX3* | TOX high mobility group box family member 3 |
| *TPBG* | trophoblast glycoprotein |
| *TPBG* | trophoblast glycoprotein |
| *TPT1* | tumor protein, translationally-controlled 1 |
| *TRIB2* | tribbles pseudokinase 2 |
| *TRIM16* | tripartite motif containing 16 |
| *TRIM24* | tripartite motif containing 24 |
| *TRIM59* | tripartite motif containing 59 |
| *TRIM67* | tripartite motif containing 67 |
| *TRIO* | trio Rho guanine nucleotide exchange factor |
| *TRIP10* | thyroid hormone receptor interactor 10 |
| *TTC28* | tetratricopeptide repeat domain 28 |
| *TTC30B* | tetratricopeptide repeat domain 30B |
| *TTC5* | tetratricopeptide repeat domain 5 |
| *TTK* | TTK protein kinase |
| *TUBA1A* | tubulin alpha 1a |
| *TULP3* | tubby like protein 3 |
| *TXNIP* | thioredoxin interacting protein |
| *TYMS* | thymidylate synthetase |
| *UBE2J1* | ubiquitin conjugating enzyme E2 J1 |
| *UBE2L6* | ubiquitin conjugating enzyme E2 L6 |
| *UBTD2* | ubiquitin domain containing 2 |
| *UGDH* | UDP-glucose 6-dehydrogenase |
| *UHRF1* | ubiquitin like with PHD and ring finger domains 1 |
| *UIMC1* | ubiquitin interaction motif containing 1 |
| *UMPS* | uridine monophosphate synthetase |
| *USP1* | ubiquitin specific peptidase 1 |
| *USP37* | ubiquitin specific peptidase 37 |
| *USP42* | ubiquitin specific peptidase 42 |
| *USP6NL* | USP6 N-terminal like |
| *VASP* | vasodilator stimulated phosphoprotein |
| *VCAM1* | vascular cell adhesion molecule 1 |
| *VEPH1* | ventricular zone expressed PH domain containing 1 |
| *WAPL* | WAPL cohesin release factor |
| *WDR3* | WD repeat domain 3 |
| *WIZ* | widely interspaced zinc finger motifs |
| *WSCD1* | WSC domain containing 1 |
| *XPNPEP1* | X-prolyl aminopeptidase 1 |
| *XRCC1* | X-ray repair cross complementing 1 |
| *XRN2* | 5'-3' exoribonuclease 2 |
| *YLPM1* | YLP motif containing 1 |
| *YTHDF2* | YTH N6-methyladenosine RNA binding protein 2 |
| *ZBED9* | zinc finger BED-type containing 9 |
| *ZBTB14* | zinc finger and BTB domain containing 14 |
| *ZBTB34* | zinc finger and BTB domain containing 34 |
| *ZBTB46* | zinc finger and BTB domain containing 46 |
| *ZBTB6* | zinc finger and BTB domain containing 6 |
| *ZCCHC3* | zinc finger CCHC-type containing 3 |
| *ZFP36L1* | ZFP36 ring finger protein like 1 |
| *ZFP36L2* | ZFP36 ring finger protein like 2 |
| *ZFP37* | ZFP37 zinc finger protein |
| *ZFPM2* | zinc finger protein, FOG family member 2 |
| *ZKSCAN2* | zinc finger with KRAB and SCAN domains 2 |
| *ZMIZ1* | zinc finger MIZ-type containing 1 |
| *ZMYM2* | zinc finger MYM-type containing 2 |
| *ZNRD1* | zinc ribbon domain containing 1 |
| *ZNRF1* | zinc and ring finger 1 |
| *ZWILCH* | zwilch kinetochore protein |

Supplementary Figure 1: Both ePRS-IR based on ADHD GWAS 2010 (Neale, Medland et al. 2010) or ADHD GWAS 2019 (Demontis, Walters et al. 2019) are capable of predicting IST performance in boys, pre-clumping.

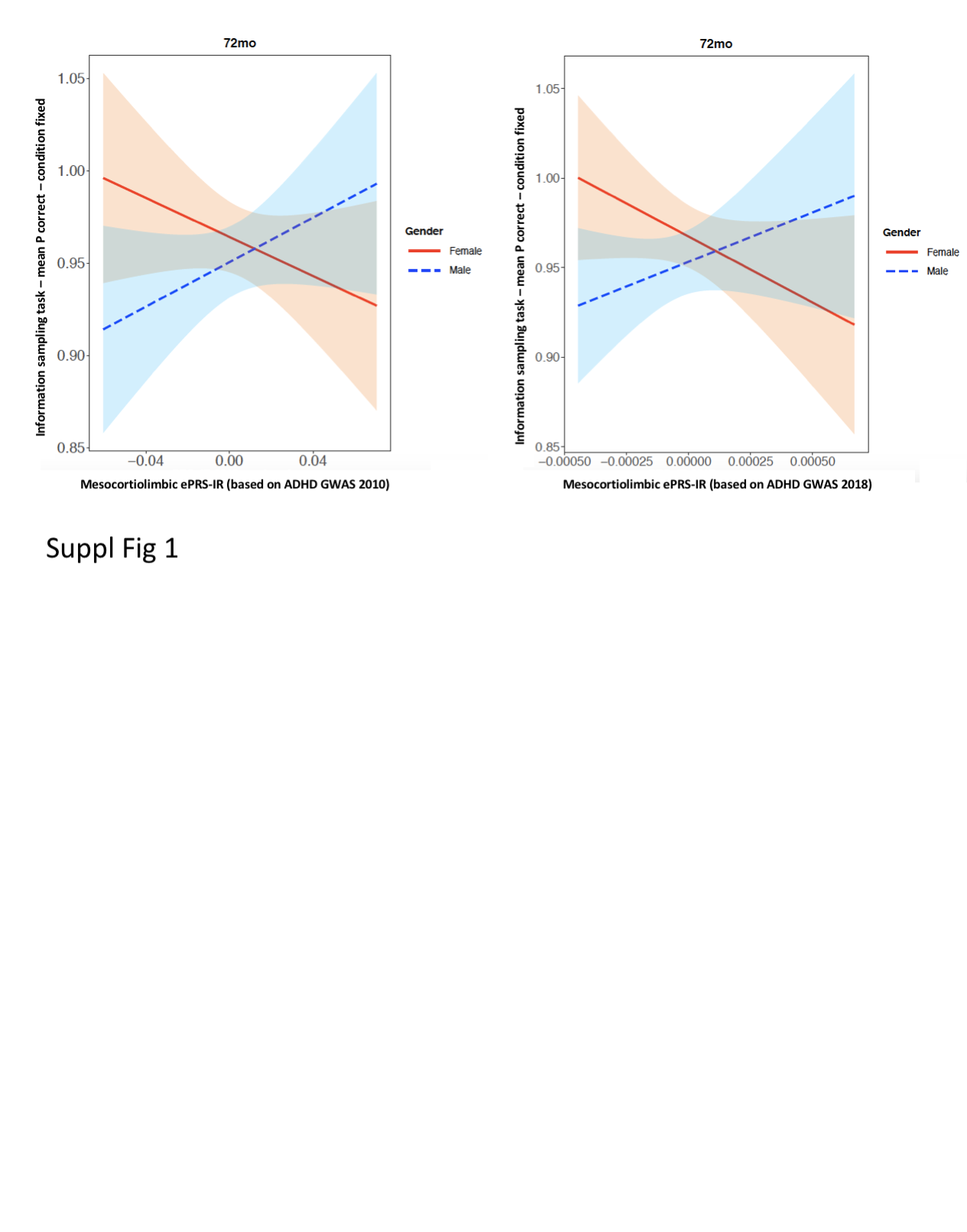
